# The role of the rodent lateral orbitofrontal cortex in simple Pavlovian cue-outcome learning depends on training experience

**DOI:** 10.1101/2020.10.16.342832

**Authors:** Marios C. Panayi, Simon Killcross

## Abstract

The orbitofrontal cortex (OFC) is a critical structure in the flexible control of value-based behaviours. OFC dysfunction is typically only detected when task or environmental contingencies change, against a backdrop of apparently intact initial acquisition and behaviour. While intact acquisition following OFC lesions in simple Pavlovian cue-outcome conditioning is often predicted by models of OFC function, this predicted null effect has not been thoroughly investigated. Here we test the effects of lesions and temporary muscimol inactivation of the rodent lateral OFC on the acquisition of a simple single cue-outcome relationship. Surprisingly, pre-training lesions significantly enhanced acquisition after over-training whereas post-training lesions and inactivation significantly impaired acquisition. This impaired acquisition to the cue reflects a disruption of behavioural control and not learning since the cue could also act as an effective blocking stimulus in an associative blocking procedure. These findings suggest that even simple cue-outcome representations acquired in the absence of OFC function are impoverished. Therefore, while OFC function is often associated with flexible behavioural control in complex environments, it is also involved in very simple Pavlovian acquisition where complex cue-outcome relationships are irrelevant to task performance.

## Introduction

The orbitofrontal cortex (OFC) is critical to behavioural flexibility when learning and behaviour need to be updated to reflect a change in the environment (Gardner et al., 2019; Klein-Flugge et al., 2013; Kringelbach, 2005; Murray & Rudebeck, 2018; Rudebeck & Murray, 2014). In particular, the OFC is necessary for appropriately updating behaviour when the contingencies between predictive cues and outcomes change, or when outcomes change in value (Panayi & Killcross, 2018; Pickens et al., 2005; Walton et al., 2011).The information encoded in OFC about the expected value and identity of predicted outcomes is necessary for flexibly updating behaviour when these outcome features change. Population and single-unit neuronal firing in the OFC encodes many features of reward outcomes (e.g. size, preference, identity, time, location, probability, certainty, salience (Delamater, 2007; Ogawa et al., 2013; Padoa-Schioppa, 2009; Sadacca et al., 2018; Stalnaker et al., 2014; Takahashi et al., 2013; Zhou et al., 2019)), furthermore the coding of these features develops over the course of learning to predictive cues in anticipation of the expected outcome (Schoenbaum et al., 2009). There is also substantial evidence to suggest that this outcome expectancy information in the OFC is incorporated into mid-brain dopaminergic reward prediction errors (Takahashi et al., 2011), which are critical for learning (Schultz, 1998; Steinberg et al., 2013).

However, despite these close ties to the learning process, the OFC is typically not necessary for the initial learning (comprehensively documented by Delamater, 2007; Izquierdo, 2017; Murray et al., 2007; Rudebeck & Murray, 2014; Stalnaker et al., 2015), except in the most complex of circumstances (e.g. Walton et al., 2011). Lesions and functional inactivation of the OFC do not appear to disturb initial learning about Pavlovian cue-outcome relationships in a range of tasks, and instead only reveal their effects when the cue-outcome relationships change, or when the value of expected outcomes change, such as in reversal learning and outcome devaluation procedures (Butter, 1969; Dias et al., 1996; Gallagher et al., 1999; Iversen & Mishkin, 1970; Schoenbaum et al., 2003; West et al., 2011). To account for these effects, one class of OFC theories suggests that the OFC is necessary for representing information about the sensory-specific properties or identity of expected outcomes (Burke et al., 2008; Delamater, 2007; Schoenbaum et al., 2009, 2011). A second, but complementary class of theories using a reinforcement learning framework suggests that the OFC is necessary for the representation of latent state information (Wilson et al., 2014). In reinforcement learning models, tasks such as Pavlovian conditioning can be divided into discrete physically observable states, such as “cue on”, “cue off”, and “reward”, and underlying latent states signalled by partially observable information recalled into working memory such as reinforcement history.

Both theories, while couched in different computational and theoretical frameworks, suggest similar roles for the OFC. Latent states encompass specific outcome expectancies and include a broader category of potential stimuli (e.g. internal context (Niv, 2019)). Implicit in these theories is that initial acquisition should be affected by OFC dysfunction if performance depends on specific outcome expectancy or latent states (e.g. the differential outcomes effect (Boulougouris et al., 2007; Boulougouris & Robbins, 2009; McDannald et al., 2005); complex multiple-choice probabilistic learning tasks (Walton et al., 2011)), but not in putatively “simple” single CS-US learning tasks (Gallagher et al., 1999) where the outcome identity and value of the US stays constant and is reliably predicted by the CS. While this null effect is often reported in procedures involving learning about multiple CSs and/or USs (Burke et al., 2008; Panayi & Killcross, 2018; Schoenbaum et al., 2009), there is little evidence from tasks involving only a single CS-US relationship where a null result is clearly predicted. For example, Gallagher et al (1999) found no effect of complete OFC lesions on single CS-US acquisition but stopped training before behaviour reached asymptote (Schoenbaum et al., 2003).

Both latent state and sensory-specific outcome expectancy theories of OFC function predict a null effect of OFC lesions on initial acquisition learning, particularly in situations involving only a single CS-US relationship. Indeed, this null effect is often reported as an important feature of OFC dysfunction as it demonstrates that behaviour can appear normal when the impoverished aspects of the underlying task representation are not directly relevant to task performance (Murray et al., 2007; Schoenbaum et al., 2009; Stalnaker et al., 2015; Wilson et al., 2014). Here we directly tested this prediction in rats trained on a single CS-US Pavlovian task following lesions targeting the lateral OFC. Surprisingly, pre-training OFC lesions significantly increased Pavlovian acquisition behaviour after extended training. In contrast, post-training lesions and intra-OFC infusions of muscimol impaired Pavlovian acquisition behaviour. Using an associative blocking design, we confirmed that even though behaviour was impaired, the underlying learning about the CS-US contingency was left intact.

## Results

### Experiment 1: Pre-training OFC lesions

#### Acquisition

Pre-training OFC lesions significantly increased responding to the Pavlovian cue relative to sham control animals (Figure 1A; lesions depicted in Figure 1-figure supplement 1). Analysis of conditioned responding was conducted as a CS-PreCS difference score such that levels of responding reflected discriminative performance to the cue (CS) above baseline (PreCS). Acquisition of responding to the CS was significantly greater in the lesion group than the sham group (main effect of Group *F*(1,40) = 10.83, *p* = .002, Block *F*(6,240) = 34.07, *p* < .001, and Group x Block interaction *F*(6,240) = 7.33, *p* < .001). Follow up comparisons on each block revealed that responding in the lesion group was significantly higher than the sham group during the last 4 blocks (Block 1 *t*(40) = −1.67, *p* = .103, Block 2 *t*(40) = 0.14, *p* = .893, Block 3 *t*(40) = 1.79, *p* = .082, Block 4 *t*(40) = 2.39, *p* = .022, Block 5 *t*(40) = 4.59, *p* < .001, Block 6 *t*(40) = 3.48, *p* = .001, Block 7 *t*(40) = 2.32, *p* = .026). Given the ubiquity of non-significant effects of OFC lesions on acquisition learning in the literature, two independent replications of this novel effect were conducted (combined here; same pattern of statistical significance in both independent replications) which confirmed the effect was robust.

**Figure 1.**
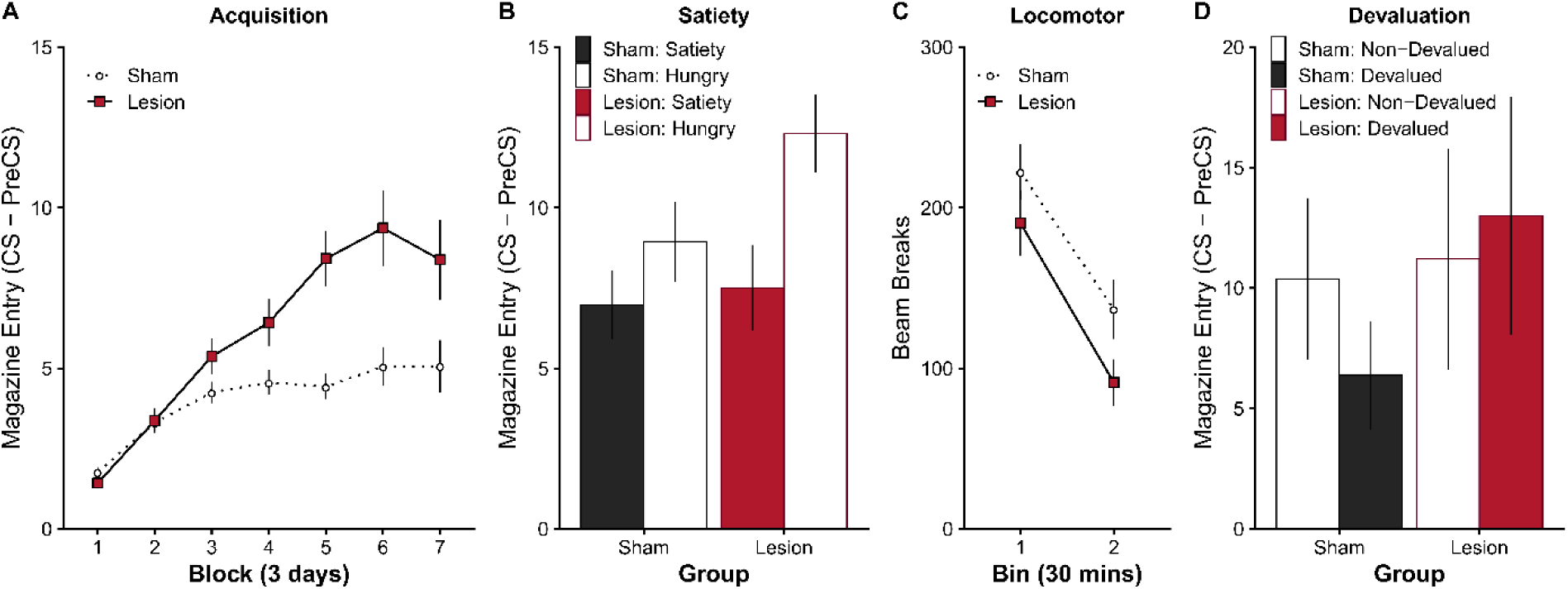
The effect of pre-training OFC lesions on the acquisition of simple Pavlovian cue-outcome relationship. Representative lesion damage and histology depicted in <u>Figure 1-figure supplement 1</u>. (**A**) Experiment 1: OFC lesions significantly enhance acquisition behaviour to a simple Pavlovian cue (CS) predicting a food pellet. Responding during the baseline PreCS period is subtracted from responding during the CS period (i.e. CS-PreCS). Data presented in blocks of 3 days. (**B**) The effect of manipulating general levels of satiety (24 hrs ad-libitum access to food) on Pavlovian acquisition behaviour in a subset of rats (subgroup 1; sham n = 8, lesion n = 7). General satiety reduced behaviour in the lesion group and abolished group differences (sated), which returned when tested hungry 24 hours later. The effect of satiety was also evident on the first trial of the session <u>Figure 1-figure supplement 2</u>. (**C**) Locomotor activity (as reflected by infra-red beam breaks in a novel open-field) measured over 1 hour (separated into 30 min blocks) shows no significant hyperactivity in the OFC lesion group. (**D**) The effect of outcome-specific devaluation is abolished by OFC lesions (subgroup 2; sham *n* = 8, lesion *n* = 5). After retraining with two unique Pavlovian cues and outcomes (<u>Figure 1-figure supplement 3A</u>), one outcome was paired with injections of LiCl to establish an outcome specific taste aversion (<u>Figure 1-figure supplement 3B</u>). At test, responding to the cue that predicted the now Devalued outcome (vs the Non-Devalued control outcome) revealed that the sham group appropriately reduced behaviour for the Devalued outcome whereas the lesion group did not. Error bars depict ± SEM.

#### Locomotor activity

The enhanced responding observed during acquisition in the OFC lesion group could simply reflect an enhancement of general locomotor activity. However locomotor activity (Figure 1C) did not differ between groups (main effect of TimeBin *F*(1,33) = 62.93, *p* < .001, but no significant effect of Group *F*(1,33) = 2.87, *p* = .100, or Group x TimeBin interaction *F*(1,33) = 0.36, *p* = .555). Therefore, the enhanced responding during acquisition was not simply due to lateral OFC lesions inducing hyperactivity, consistent with previous findings (e.g. Lasseter et al., 2009; Panayi & Killcross, 2018).

#### General Satiety

To test whether the enhanced responding following pre-training OFC lesions was sensitive to levels of hunger or shifts in general motivation, a subgroup of animals (subgroup 1) was tested when sated, i.e. following 24 hours *ad libitum* access to home-cage food (Figure 1B). General satiety, did not affect the rate of responding in the sham group (Sham: Satiety vs Hungry *t*(13) = −1.38, *p* = .191) but significantly suppressed responding in the lesion group (Lesion: Satiety vs Hungry *t*(13) = −4.24, *p* = .001) compared to subsequent testing 24 hours later when hungry again (no significant main effect of Group *F*(1,13) = 1.43, *p* = .253, but a significant main effect of Hunger *F*(1,13) = 16.30, *p* = .001, and Group x Hunger interaction *F*(1,13) = 4.63, *p* = .051). Since the satiety test session was rewarded, it is possible that OFC lesioned animals could learn that the reward was less valuable by direct experience with the reward, similar to incentive learning effects normally observed in instrumental conditioning (Dickinson & Balleine, 2002). However, this possibility is unlikely as responding was comparable between groups on the first trial of the satiety test (*t*(13) = 1.04, *p* = .317, Figure 1-figure supplement 2), before the first reward was delivered. This suggests that, consistent with previous reports (e.g. McDannald et al., 2005), animals with lateral OFC lesions are sensitive to shifts in hunger and general motivation.

#### Devaluation Test

OFC lesions have been shown to cause characteristic deficits in Pavlovian outcome devaluation (Gallagher et al., 1999; Panayi & Killcross, 2018; Pickens et al., 2003, 2005). Therefore, to test whether the present lesion manipulation was comparable to other reports we tested a subgroup of animals (subgroup 2) on Pavlovian outcome devaluation. First the sham and lesion animals were given novel acquisition training of two novel and unique cue-outcome relationship (Figure 1-figure supplement 3A). A specific taste aversion was then established by pairing consumption of one of the outcomes with illness (i.p. injection of lithium chloride; Devalued), and the value of the other outcome was left intact (i.p. injection of saline; Non-Devalued). Both groups learned the novel cue-outcome associations and acquired the specific taste aversion (Figure 1-figure supplement 3B).

Finally, during a devaluation test (Figure 1D), the two cues were presented in extinction. The sham group showed a significant devaluation effect, i.e. responding was lower to the devalued than non-devalued cue (*t*(11) = −3.06, *p* = .011). In contrast, the devaluation effect was abolished in the lesion group, and responding remained high to both the devalued and non-devalued cue (*t*(11) = 1.09, *p* = .300; Significant Group x Cue interaction *F*(1,11) = 7.55, *p* = .019, but no main effect of Group *F*(1,11) = 0.54, *p* = .479, or Cue *F*(1,11) = 1.09, *p* = .320). This finding successfully replicates the finding that both complete OFC and focal lateral OFC lesions abolish the outcome devaluation effect in rodents (Gallagher et al., 1999; Panayi & Killcross, 2018; Pickens et al., 2003, 2005).

### Experiment 2: Post-training muscimol inactivation

The enhanced Pavlovian responding observed following OFC lesions (Figure 1A) may be due to enhanced learning of a general cue-outcome predictive relationship in the OFC lesion group (Figure 2-figure supplement 1). This is consistent with a role for the OFC in representing outcome expectancy information. For example, incremental learning about a cue-outcome relationship is thought to depend upon prediction errors (Esber & Haselgrove, 2011; LePelley, 2004; Mackintosh, 1975; Nasser et al., 2017; Pearce & Hall, 1980; Rescorla & Wagner, 1972; Sutton & Barto, 1998), i.e. the difference between the experience outcome value and the expected outcome value. The expected outcome value of a cue is incrementally updated until this prediction error discrepancy is minimised. If the OFC carries some aspect of outcome expectancy information (Baxter et al., 2000; Pears et al., 2003; Schoenbaum et al., 2009; Takahashi et al., 2009, 2011), then OFC lesions might consistently reduce/underestimate the expected value of a cue which in turn would result in abnormally persistent prediction errors and enhanced learning. Therefore, disruption of OFC function should temporarily lower expected value, and enhance prediction errors and learning supported by other brain regions (for modelling of this prediction see Figure 2-figure supplement 1). We tested this hypothesis by inactivating the OFC after first successfully acquiring cue-outcome learning i.e. when expected value is high and prediction errors are low. If the OFC carries some aspect of the learned expected value, then inactivation of the OFC should restore prediction errors at the time of reward and responding should increase to reflect new learning. Following this, returning function to the OFC should result in an over-expectation of the value of the outcome, and performance should decrease to reflect the extinction of this over-expectation. Importantly, while this account is couched in terms prediction-error learning mechanisms, the prediction remains true for any account of OFC lesions enhancing learning (Figure 2-figure supplement 1).

We tested this hypothesis by first training a new group of animals on the same simple Pavlovian task for 9 days, before implantation of bilateral cannulae targeting the OFC (Figure 2, Days 1-9; significant main effect of Day *F*(8,176) = 25.42, *r* < .001, but no main effect of Group *F*(1,22) = 1.08, *r* = .310, or Group x Day interaction *F*(8,176) = 0.54, *r* = .825). Following post-operative recovery (histology depicted in Figure 2-figure supplement 2), and prior to infusion, response levels were similar in both groups (Figure 2, Post; no significant differences between Groups *t*(22) = −0.68, *r* = .501).

**Figure 2.**
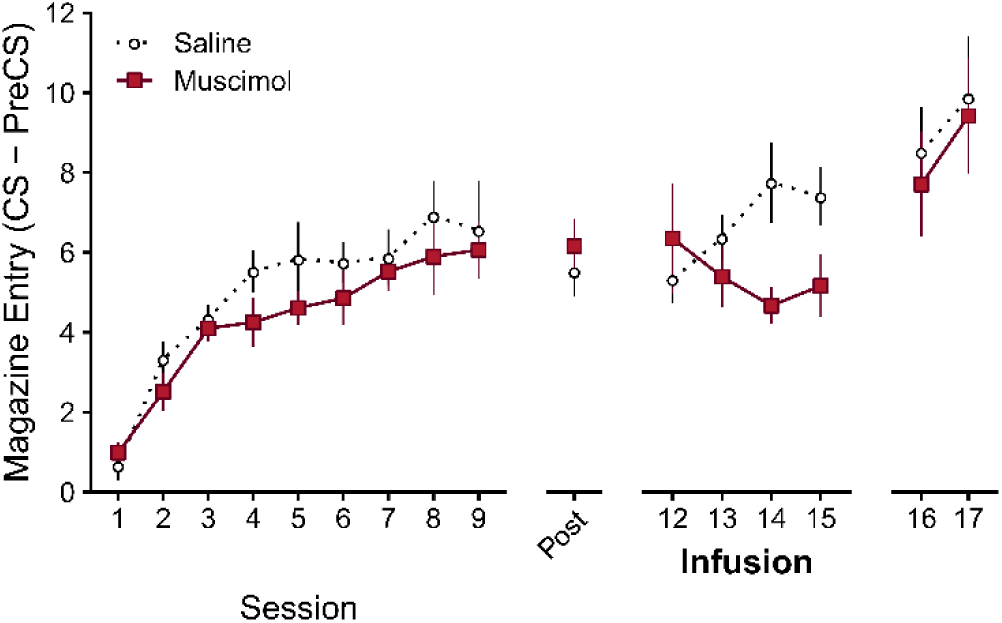
Post-training OFC inactivation suppresses Pavlovian acquisition behaviour, in contrast to pre-training lesions which increased Pavlovian acquisition behaviour. If pre-training lesions increase Pavlovian learning, then post-training lesions or inactivation should also increase learning (rationale and learning model predictions in <u>Figure 2-figure supplement 1</u>). Experiment 2: Rates of discriminative responding (CS-PreCS) during initial acquisition (sessions 1-9), post-operative recovery (post), following intra-OFC infusion of muscimol or saline (sessions 12-15), and without infusion (sessions 16-17). Cannulae placements depicted in <u>Figure 2-figure supplement 2</u>. The effect of post-training lesions on acquisition revealed the same pattern of results (<u>Figure 2-figure supplement 3, Figure 2-figure supplement 4</u>). Error bars depict ± SEM.

Contrary to our prediction, intra-OFC muscimol infusions disrupted rather than enhanced further acquisition of responding relative to the saline group (Figure 2, Infusion - Days 12-15; Significant Group x Day interaction *F*(3,66) = 5.03, *p* = .003, but no main effect of Group *F*(1,22) = 1.90, *p* = .182, or Day *F*(3,66) = 0.32, *p* = .809). Simple effects revealed significantly greater responding in the saline group on the last 2 days of infusions (Muscimol vs Saline: Day 12 *t*(22) = 0.67, *p* = .508, Day 13 *t*(22) = −1.03, *p* = .315, Day 14 *t*(22) = −2.79, *p* = .011, Day 15 *t*(22) = −2.08, *p* = .049). Furthermore, the saline group increased responding across infusion days 12-15 (Saline: significant positive linear trend *t*(22) = 2.79, *p* = .011), whereas the muscimol group did not (Muscimol: no significant linear trend *t*(22) = −1.57, *p* = .131). Therefore, post-training inactivation of the OFC impaired acquisition.

Post-infusion, with function returned to the OFC, the group differences observed under drug infusion were no longer apparent, and both groups continued to acquire responding at similar levels (Figure 2, Days 16-17; significant main effect of Day *F*(1,22) = 16.05, *p* = .001, but no main effect of Group *F*(1,22) = 0.11, *p* = .740, or Group x Day interaction *F*(1,22) = 0.21, *p* = .649). Therefore, the effect of OFC inactivation did not persist, which suggests that the disruption in acquisition following OFC inactivation might not have impaired learning *per se*.

Furthermore, we tested post-training lesions to rule out the possibility that the differences between pre- and post-training OFC manipulations were simply due to differences in the method of manipulation i.e. excitotoxic lesions vs inactivation using a GABA-A agonist. Consistent with muscimol inactivation, post-training lesions significantly impaired Pavlovian acquisition (Supplementary Experiment 1: Figure 2-figure supplement 3, Figure 2-figure supplement 4). Therefore, it is unlikely that the difference between pre- and post-training OFC manipulations observed in Experiment 1 and 2 are due to the method of manipulation.

### Experiment 3: OFC inactivation prior to associative blocking

OFC inactivation during acquisition suppressed cue responding, but it is unclear if this reduction in behaviour is due to suppression of additional learning or behavioural performance (Figure 2). This ambiguity is predominantly driven by the assumption that an animal’s response levels represent some monotonic function of acquired learning (Mackintosh, 1975; Pearce & Hall, 1980; Rescorla & Wagner, 1972; Sutton & Barto, 1998; Wagner, 1981). To disambiguate learning from performance effects we employed an associative blocking design (Figure 3A). In a blocking experiment, first an animal is trained such that a cue (cue A) predicts an outcome (pellet). Next, A is presented in compound with a novel cue (cue B) which also leads to the same pellet outcome. If the animal has learned that cue A sufficiently predicts the pellet outcome already, then very little is learned about cue B i.e. learning about cue A blocks subsequent learning about cue B (Kamin, 1969). However, if learning about cue A is insufficient, then learning about cue B should not be blocked. We predicted that if OFC inactivation is disrupting learning, then OFC inactivation during initial learning about cue A should disrupt the blocking effect.

**Figure 3.**
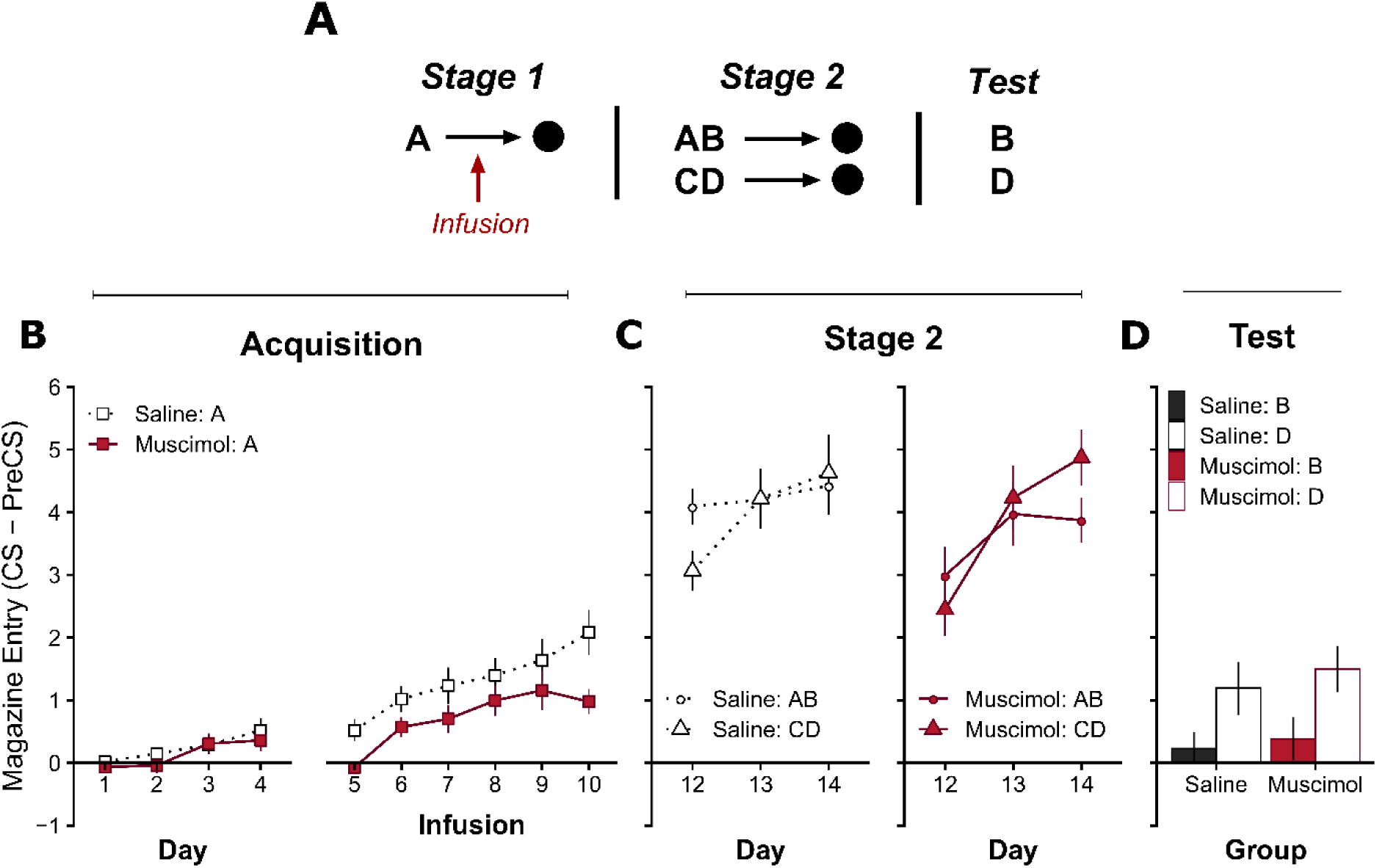
The effect of OFC inactivation during acquisition on subsequent learning in a Pavlovian blocking design. Experiment 3: (A) The design used to achieve blocking of learning to cue B during stage 2 by pre-training cue A in stage 1. OFC infusions of saline or muscimol were performed during stage 1 after the first 4 days of initial acquisition to cue A. Cues A and C were always visual cues, either darkness caused by extinguishing the houselight or flashing panel lights (5Hz). Cues B and D were always auditory cues, either an 80dB white noise or a 5Hz train of clicks. All cues lasted 10s, and reward was always a single food pellet. Cannulae placements depicted in Figure 3-figure supplement 3. (B) Pavlovian acquisition to cue A over 10 days, with intact OFC (days 1-4) and following infusion of saline or muscimol to functionally inactivate the OFC (days 5-10). Muscimol infusions significantly suppressed responding to cue A. (C) Performance during stage 2 of blocking to cue compounds AB and CD in the saline (left) and muscimol (right) infusion groups. A focused analysis of responding within Day 12 is presented in Figure 3-figure supplement 4. (D) Responding during an extinction test to “blocked” cue B and the overshadowing control cue D. Figure 3-figure supplement 5 shows subsequent reacquisition to cues B and A to assess possible differences in attentional strategies between the saline and muscimol group. Significantly reduced responding to cue B relative to cue D indicates that learning about cue A effectively blocked subsequent learning to cue B in both the muscimol and saline groups. Pavlovian responding quantified by the rate of discriminative responding (CS-PreCS). Error bars depict ± SEM.

First, we demonstrated again that OFC inactivation significantly impairs acquisition in a new cohort of animals using similar parameters to those required for the associative blocking design (inactivation from days 5-10 of acquisition with a 10s visual CS; Supplementary Experiment 2: Figure 3-figure supplement 1, Figure 3-figure supplement 2). Again, OFC inactivation significantly impaired acquisition, confirming that the effects observed in Figure 2 are not dependent on a specific cue modality or duration.

Next, in a different cohort of animals, we tested whether impaired CS-US acquisition following OFC inactivation disrupted subsequent Pavlovian blocking (cannula placements depicted in Figure 3-figure supplement 3). During stage 1 of blocking (Figure 3B), all animals were given 10 days of acquisition training to cue A. OFC function was intact during the first 4 days of acquisition, and all animals began to acquire the cue A-outcome relationship (Days 1-4: significant main effect of Day *F*(3,72) = 5.77, *p* = .001, but no effect of Group , or Group x Day interaction *F*(3,72) = 0.27, *p* = .850). All animals then received an additional 6 days of acquisition to cue A (Figure 3B, Days 5-10) following either intra-OFC infusions of muscimol or saline. Infusions of muscimol depressed overall responding relative to saline infusions (significant main effect of Group *F*(1,24) = 4.25, *p* = .050, and Day *F*(5,120) = 17.49, *p* < .001, but no Group x Day interaction *F*(5,120) = 1.31, *p* = .263). Importantly, on the final day (Day 10), responding in the muscimol group was significantly lower than the saline group (*t*(24) = −2.69, *p* = .013).

Next, animals were trained such that compounds AB and CD also predicted reward (Figure 3C, Stage 2), importantly OFC function was intact in all animals i.e. no infusions. Responding in both the saline and muscimol groups was initially lower to the novel compound CD than to AB (Significant Cue x Day interaction *F*(2,48) = 12.12, *p* < .001, and main effect of Day *F*(2,48) = 20.09, *p* < .001, but no other main effects or interactions with Group were significant, all remaining effects *F* < 1.91, *p* > .160; Cue AB vs CD: Day 12 *t*(24) = 3.74, *p* = .001, Day 13 *t*(24) = −0.44, *p* = .663, Day 14 *t*(24) = −1.80, *p* = .085). However, the pattern of means suggests that responding to compound AB in the muscimol group was similar to the novel compound CD on Day 12 (Figure 3C, Right - Day 12, Muscimol: AB vs CD *t*(24) = 1.82, *p* = .081), and lower than compound AB in the saline group (Figure 3C, Left - Day 12; Day 12, Saline: AB vs CD *t*(24) = 3.47, *p* = .002). Furthermore, Within-session changes over trials on Day 12 revealed rapid within-session acquisition to both compounds in both groups, but responding was significantly lower in the muscimol group at the start of the session (Figure 3 - figure supplement 4; First 2 trials, significant main effect of Group *F*(1,24) = 8.67, *p* = .007, and Cue *F*(1,24) = 7.61, *p* = .011, but no Group x Cue interaction *F*(1,24) = 0.19, *p* = .670). The lower responding to cue AB in the muscimol group suggests that acquisition to cue A was impaired following infusions in Stage 1 and this impairment persisted (albeit transiently) when test drug free in stage 2. Indeed, the levels of responding to compound AB in the muscimol group at the start of Day 12 (Figure 3 - figure supplement 4) are similar to levels of responding to the novel compound CD in the saline group. This would suggest that learning about cue A in the muscimol group was impaired in stage 1, and therefore cue A should not effectively block learning to cue B in stage 2.

At test both groups showed significant blocking of learning to cue B relative to the control cue D (Figure 3D; Significant main effect of Cue *F*(1,24) = 7.29, *p* = .013, but no main effect of Group *F*(1,24) = 0.54, *p* = .471, or Group x Cue interaction *F*(1,24) = 0.04, *p* = .843). This suggests that inactivation of the OFC significantly reduced behavioural performance but not learning to cue A in Stage 1, and this impairment transiently affected compound AB on Day 12 in the absence of OFC inactivation. Therefore, the impairments observed in our earlier findings (Figure 2A, post infusion) are unlikely to be due to impairments in learning. In addition to this, we rule out the possibility that the two groups used different attentional solutions to achieve a similar blocking result (Figure 3-figure supplement 5).

## Discussion

The present studies tested the hypothesis that the rodent lateral OFC is not necessary for Pavlovian acquisition in simple single CS-US procedure. Here we show that OFC lesions and inactivation significantly affects Pavlovian acquisition. Furthermore, we found a dissociation between pre- and post-training OFC manipulations on Pavlovian acquisition such that pre-training OFC lesions enhance, whereas post-training lesions and inactivation impairs acquisition behaviour. Given the absence of these effects in the extant literature, it is notable that these effects were robust and were replicated multiple times. Next, using an associative blocking design, we tested whether impaired behaviour following post-training OFC inactivation reflects a disruption of learning or behavioural control. OFC inactivation did not disrupt the underlying learning about the predictive CS-US relationship as assayed by blocking, and instead disrupted the appropriate control of anticipatory behaviour to the CS.

### Lateral OFC is necessary for simple Pavlovian acquisition

The significant role of the OFC in Pavlovian acquisition in the present studies is surprising since OFC lesions and inactivation have consistently been reported to have no effect on acquisition in rats (e.g. Burke et al., 2008; Gallagher et al., 1999; Schoenbaum et al., 2002; Stalnaker et al., 2007), unless there are complex cue- or outcome-specific task demands (e.g. Ramirez & Savage, 2007). However, in tasks involving simple single Pavlovian CS-US procedures and pre-training OFC lesions, performance often does not reach asymptote (e.g. after 9 days, (Gallagher et al., 1999)) before proceeding to a new stage of the experiment. In Experiment 1, we did not observe any significant effects of OFC lesions on acquisition until around 15-21 days of acquisition. However, after extended training Schoenbaum et al (2003) have reported significant effects of OFC lesions on acquisition in a simple cue-outcome go-nogo task when looking at response latencies, but not on trials-to-criterion. Therefore, the effects of pretraining lesions may not have been observed previously due to task specific parameters such as the length of training and the sensitivity of the response measures.

Pretraining OFC lesions have been shown to disrupt Pavlovian acquisition in sign-tracking procedures in which lever insertion is used as the CS (Chudasama & Robbins, 2003). Focal lateral OFC lesions also significantly impair sign-tracking behaviour (i.e. engaging with the lever cue), and bias behaviour towards goal-tacking (i.e. approaching the magazine) (Panayi & Killcross, 2018). The present findings that pre-training OFC lesions enhanced behaviour focused toward the magazine is consistent with a deficit in sign-tracking and a bias towards goal-tracking.

In contrast to pre-training lesions, post-training OFC inactivation/lesions normally coincide with changes in experimental phase and continued acquisition is not assessed. In tasks in which OFC inactivation coincides with a change in experimental stage, the effects of OFC inactivation are consistent with an impairment in subsequent acquisition (2009). For example, Burke et al (2009) found that post-training OFC inactivation impaired acquisition to a Pavlovian CS in reversal task. Similarly, Takahashi et al (2009) found that OFC inactivation during a Pavlovian over-expectation task disrupted new learning. Therefore, the robust effect of impaired acquisition following post-training OFC inactivation that we report is consistent with impaired subsequent acquisition in tasks with more complex manipulations.

### Lateral OFC is not necessary for learning the predictive CS-US relationship

Post-training OFC inactivation significantly impaired acquisition behaviour (Experiment 2), and this disruption was more profound when inactivation occurred earlier in training and more likely to persist after OFC function returned (Supplementary Experiment 2). This seems to suggest that learning about the CS-US relationship was disrupted. The idea that the OFC could be involved in learning is also consistent with a role for the OFC in the representation of expected values (Burke et al., 2008; Schoenbaum et al., 2011; Stalnaker et al., 2018) which influence mid-brain dopaminergic prediction errors (Takahashi et al., 2009, 2011), known to be necessary for Pavlovian learning (Sharpe et al., 2017a; Steinberg et al., 2013).

Unexpectedly, the impaired acquisition we observed following post-training OFC disruption did not disrupt the ability of the CS to block learning about a novel cue (Experiment 3, Figure 3), despite significantly impaired performance post-inactivation (Figure 3-figure supplement 4; Muscimol AB is as low as Saline CD which does not show evidence of blocking). This is surprising given that in some Pavlovian learning contexts, levels of behavioural expression can dictate the extent to which learning occurs (Delamater, 2004). This finding highlights the importance of using multiple tests of learning (Rescorla, 2002a, 2002b) to assess disrupted acquisition effects.

Intact blocking despite impaired acquisition behaviour suggests that OFC inactivation did not disrupt the underlying learning about the associative strength of the CS-US relationship. Associative blocking is often used to assess the role of prediction-error based learning (Sharpe et al., 2017b; e.g. Steinberg et al., 2013), suggesting that the OFC is not necessary for this aspect of Pavlovian learning. This distinction suggests that the learned value of a Pavlovian CS-US association might be independent of the current expected or subjective value of expected reward. Informally, learning whether an outcome will be delivered might reasonably be separate from learning the subjective value or identity of that outcome (Delamater, 2007; Delamater & Oakeshott, 2007; McDannald et al., 2011; Zhou et al., 2019).

### Pre- vs post-training effects

The dissociable and opposite effects of pre- and post-training OFC lesions/inactivation on acquisition were surprising and rule out a simple account of OFC dysfunction in terms of prediction-error based learning impairments (Figure 2-figure supplement 1). One possibility is that pretraining lesions result in compensatory function such that learning is supported by other neural systems. In contrast, post-training lesions and inactivation disrupts learning/behaviour that has been acquired in an OFC dependent manner. This argument has been proposed when only pre-training OFC lesions (Boulougouris et al., 2007; Boulougouris & Robbins, 2009), or only post-training OFC lesions disrupt behaviour (Balleine et al., 2011; Ostlund & Balleine, 2007). We will also consider two alternative accounts of pre- vs post-training OFC lesion differences based on theoretical accounts of OFC function, sensory-specific outcome expectancy and latent state theories. Note that these theories do not predict an effect of OFC lesions on simple Pavlovian acquisition *a priori*, and therefore require additional assumptions to account for the present data.

From an associative learning framework, even putatively “simple” single cue-outcome Pavlovian learning can involve a number of different psychological/behavioural processes (Dickinson, 1980; Hall, 2002; Holland, 1977; Konorski, 1967; Mackintosh, 1974; Rescorla, 1988). Take for example a 10s light cue that reliably predicts the delivery of a pellet reward. A rat can learn that the cue predicts the sensory-specific properties of the outcome (e.g. taste, texture, sweetness, colour, size, location etc…), or the general motivational value of that reward, or simply develop a stimulus-response habit to approach the reward location when the cue is presented. Indeed, there is experimental evidence for these multiple aspects of learning occurring during Pavlovian conditioning (for review see Delamater & Oakeshott, 2007). It is possible that pretraining OFC lesions disrupt the balance of these different aspects of Pavlovian learning and behaviour (Burke et al., 2007; Delamater, 2007).

If the OFC is necessary for the representation of the sensory specific properties of expected outcomes, then OFC lesions might allow a stimulus-response habit system to dominate behavioural control. Following pre-training lesions, this may lead to an unconstrained habit learning system (Coutureau & Killcross, 2003; Dickinson, 1985; Dolan & Dayan, 2013; Killcross & Coutureau, 2003) that is not necessarily bounded by the current value of the outcome, and overly sensitive to current general motivational states (e.g. overall hunger levels; Figure 1B) of the organism. However, once initial learning occurs with an intact OFC, the encoding of the identity of the expected outcome is likely to have occurred (e.g. Delamater & Holland, 2008). Subsequently, a post-training lesion or inactivation of the OFC is likely to affect the subsequent updating of this information. Therefore, one possible account is that the impaired acquisition behaviour we observed following post-training inactivation reflects an inability to update the current motivational value of the specific outcome that is expected.

The latent state representation account of the OFC might also be able to account for the differences observed dissociation between pre- and post-training OFC lesions on acquisition. Computational models (e.g. Wilson et al., 2014) often assume, for simplicity, that in a simple single cue-outcome procedure, the cue state (e.g. “light on”) is stable throughout acquisition. Given that the same cue is presented, and it always leads to the pellet outcome, this stable representation is a reasonable assumption. However, it is also likely that early in acquisition this state representation is not yet stable in healthy control animals (Niv, 2019). How can the animal be certain that the light cue, the testing chamber context, or the reward pellet that they see on each trial is identical to the trials they have already experienced within the session, and from previous days? The subjective experience of these states and their physical features is very likely to be different within- and between-sessions e.g. the ambient noises, odours, temperature of the context, the location and intensity of the light cue based on where the rat happens to be located when it turns on, and the gradual onset of sensory specific satiety to the pellet etc… Informally, how does the rat know that this light is the same light that they saw at the start of the session, or the day before? The perception and recognition of these states is therefore subject to differences in variables such as generalization, confidence, and certainty.

Paradoxically, in a simple and stable cue-outcome training procedure, pre-training OFC lesions may result in an accurate, but inflexible, representation of these simple task states quite rapidly. In this stable and simple training context this could lead to enhanced Pavlovian acquisition. However, in a task with multiple or uncertain cue-outcome contingencies pretraining OFC lesions might impair acquisition (Stolyarova & Izquierdo, 2017; Walton et al., 2010). However, post-training inactivation of the OFC would disrupt the ability to update already established state representations at whatever stage of certainty/stability that they have currently achieved. In the stable single cue-outcome learning situation employed in the present studies, this would result in disruption of further acquisition. Again, in a task with interference from multiple cue-outcome relationships, post-training lesions might improve performance.

### Conclusion

Here we show that the rodent lateral OFC is involved in Pavlovian acquisition learning process in an experience dependent manner. Once initial learning has taken place, the lateral OFC appears to be necessary for updating the current value of Pavlovian behaviours driven by expected outcome value. These findings raise two important issues. First, they demonstrate the importance of not interpreting a null effect of lesions on acquisition behaviour as evidence that the OFC is not involved in acquisition learning. Instead, the underlying deficit in acquisition is not being expressed or is not relevant to behavioural performance in the task yet. Second, these findings demonstrate that even within a putatively “simple” behavioural task, there are many potential underlying psychological processes that can contribute to performance and change over time. This is consistent with growing suggestions that the competition and interaction between underlying learning systems (e.g. Kool et al., 2018) is important and needs further study (Collins & Cockburn, 2020).

While the OFC has often been found not to be necessary for initial acquisition learning, recently there have been reports that simple Pavlovian acquisition is significantly impaired rather than enhanced following optogenetic inhibition of OFC function in head fixed mice (Namboodiri et al., 2019; Wang et al., 2020), in a manner that does not depend on VTA prediction error signalling. In contrast to our results, these studies target more ventral and medial OFC, which is likely to be an important anatomical distinction given the emerging evidence of functional heterogeneity within the OFC (Barreiros et al., 2020, 2021; Bradfield & Hart, 2020; Sharpe et al., 2015). Indeed, there appears to be dissociable but complementary roles of the medial and lateral OFC such that lateral OFC lesions disrupt Pavlovian whereas medial OFC lesions disrupt instrumental behavioural control (Bradfield et al., 2015, 2018; Gardner et al., 2017, 2018; McDannald et al., 2011; Ostlund & Balleine, 2007; Panayi & Killcross, 2018). This suggests that the OFC, as a whole, is engaged in the learning and flexible updating of value-based behaviours, but within the orbital subregions this process appears to be remarkably specialized for distinct types of behaviour and learning.

## Acknowledgements

We gratefully acknowledge Fred Westbrook, Nathan Holmes, David Bannerman, Mark Walton, Mehdi Khamassi, and Geoffrey Schoenbaum for their invaluable feedback. Research supported by grants awarded to Simon Killcross from the Australian Research Council (ARC Discovery Grant DP0989027 and DP120103564).

## Data availability

Data available online at: https://osf.io/tnbh7/?view_only=fe9f8762e1e54ca3be7e84e13b95c1d9

## Data availability

Raw data associated with figures are available at: https://osf.io/tnbh7/?view_only=fe9f8762e1e54ca3be7e84e13b95c1d9

## Methods and materials

### Animals

Subjects were male Long Evans rats (Monash Animal Services, Gippsland, Victoria, Australia) approximately 4 months old. Rats were housed four per cage in ventilated Plexiglass cages in a temperature regulated (22 ± 1°C) and light regulated (12h light/dark cycle, lights on at 7:00 AM) colony room. At least one week prior to behavioural testing, feeding was restricted to ensure that weight was approximately 95% of ad libitum feeding weight, and never dropped below 85%. All animal research was carried out in accordance with the National Institute of Health Guide for the Care and Use of Laboratories Animals (NIH publications No. 80-23, revised 1996) and approved by the University of New South Wales Animal Care and Ethics Committee.

### Apparatus

Behavioural testing was conducted in eight identical operant chambers (30.5 x 32.5 x 29.5 cm; Med Associates) individually housed within ventilated sound attenuating cabinets. Each chamber was fitted with a 3-W house light that was centrally located at the top of the left-hand wall. Food pellets could be delivered into a recessed magazine, centrally located at the bottom of the right-hand wall. Delivery of up to two separate liquid rewards via rubber tubing into the magazine was achieved using peristaltic pumps located above the testing chamber. The top of the magazine contained a white LED light that could serve as a visual stimulus. Access to the magazine was measured by infrared detectors at the mouth of the recess. Two retractable levers were located on either side of the magazine on the right-hand wall. A speaker located to the right of the house light could provide auditory stimuli to the chamber. In addition, a 5-Hz train of clicks produced by a heavy-duty relay placed outside the chamber at the back-right corner of the cabinet was used as an auditory stimulus. The chambers were wiped down with ethanol (80% v/v) between each session. A computer equipped with Med-PC software (Med Associates Inc., St. Albans, VT, USA) was used to control the experimental procedures and record data.

### Consumption chambers

To provide individual access to reinforcers during the satiety and devaluation procedures, rats were individually placed into an individual cage (33 x 18 x 14 cm clear Perspex cage with a wireframe top). Pellet reinforcers were presented in small glass ramekins inside the box and liquid reinforcers were presented in water bottles with a sipper tube. 1 day prior to the target procedure, all rats were exposed to the individual cages and given 30 mins of free access to home cage food and water to reduce novelty to the context and consuming from the ramekin and water bottles.

### Locomotor activity

Locomotor activity was assessed in eight identical boxes measuring 50 x 36x 18 cm (length x width x height), housed in a sound attenuated room. Each box consisted of 4 opaque white polyurethane walls and floor and a removable roof. In the center of the roof was an 18×40 cm grid of 3×3 mm ventilation holes. Two custom pairs of infrared beam detectors spanned the width of the box to detect locomotor activity and were located 15 cm from each end of the box. Beam breaks, corresponding to activity within the box, were recorded on a computer equipped with Med-PC software (Med Associates Inc.).

### Surgery

Excitotoxic lesions targeting the lateral OFC were performed in Experiment 1 and Supplementary Experiment 1. Rats were anesthetized with isoflurane, their heads shaved, and placed in a stereotaxic frame (World Precision Instruments, Inc., Sarasota, FL, USA). The scalp was incised, and the skull exposed and adjusted to flat skull position. Two small holes were drilled into the skull and the dura mater was severed to reveal the underlying cortical parenchyma. A 1-µL Hamilton needle (Hamilton Company, Reno, NV, USA) was lowered through the two holes targeting the lateral OFC (co-ordinates specified below). Stereotaxic co-ordinates were AP: +3.5 mm; ML: ±2.2 mm; D-V: -5.0 mm from bregma. At each site, the needle was first left to rest for 1 min. Then an infusion of N-methyl-D-aspartic acid (NMDA; Sigma-Aldrich, Switzerland), dissolved in phosphate buffered saline (pH 7.4) to achieve a concentration of 10μg/μL, was infused for 3 mins at a rate of 0.1 µ/min. Finally, the needle was left in situ for a further 4 mins to allow the solution to diffuse into the tissue. Following the diffusion period, the syringe was retracted, and the scalp cleaned and sutured. Sham lesions proceeded identically to excitotoxic lesions except that no drugs were infused during the infusion period. After a minimum of 1 week of postoperative recovery, rats were returned to food restriction for 2 days prior to further training.

In Experiments 2, 3, and Supplementary Experiment 2, bilateral guide cannulae were surgically implanted targeting the lateral OFC. Rats were anesthetized with isoflurane, their heads shaved, and placed in a stereotaxic frame (World Precision Instruments, Inc., Sarasota, FL, USA). The scalp was incised, and the skull exposed and adjusted to flat skull position. Two small holes were drilled for the cannulae using a high-speed drill, and four holes were hand drilled on different bone plates to hold fixing screws. Bilateral stainless steel guide cannulae (26 gauge, length 5mm below pedestal; Plastics One, Roanoke, VA, USA) were lowered into the lateral OFC (AP: +3.5 mm; ML: ±2.2 mm; D-V: -4.0 mm from bregma). Cannulae were held in place by dental cement and anchored to the skull with 4 fixing screws. Removable dummy cannulae were inserted into the guide cannulae to prevent them from blocking. After one week of postoperative recovery, rats were returned to food restriction for 2 days prior to further testing.

### Drugs and infusions

The GABA_A_ agonist muscimol (Sigma-Aldrich, Switzerland) was dissolved in 0.9% (w/v) non-pyrogenic saline to obtain a final concentration of 0.5 *μ*g/0.5 *μ*l. Non-pyrogenic saline 0.9% (w/v) was used as the saline control. During infusions, muscimol or saline was infused bilaterally into the lateral OFC by inserting a 33-gauge internal cannula into the guide cannula which extended 1 mm ventral to the guide tip. The internal cannula was connected to a 25 *μ*l glass syringe (Hamilton Company, Reno, NV, USA) attached to a microinfusion pump (World Precision Instruments, Inc., Sarasota, FL, USA). A total volume of 0.5 *μ*l was delivered to each side at a rate of 0.25 *μ*l/min. The internal cannula remained in place for an additional 1 min after the infusion and then removed. During the infusion procedure animals could move freely in a bucket to minimize stress. Dummy cannulae were removed prior to, and replaced immediately after, infusions. For the two training sessions prior to infusions, all animals received dummy infusions which were identical to the infusion procedure, except that no liquids were infused. These dummy infusions were performed to familiarize the rats with the microinfusion procedure and thereby minimize stress. Dummy infusions were also conducted on test sessions after the infusions to minimise differences in handling between experimental stages.

### Reinforcers

The reinforcers used were a single grain pellet (45 mg dustless precision grain-based pellets; Bio-serv, Frenchtown, NJ, USA), 20% w/v sucrose solution and 20% w/v maltodextrin solution (Myopure, Petersham, NSW, Australia). Liquid reinforcers were flavoured with either 0.4% v/v concentrated lemon juice (Berri, Melbourne, Victoria, Australia) or 0.2% v/v peppermint extract (Queen Fine Foods, Alderley, QLD, Australia) to provide unique sensory properties to each reinforcer. Liquids were delivered over a period of 0.33 s via a peristaltic pump which corresponded to a volume of 0.2 mL. The volume and concentration of liquid reinforcers was chosen to match the calorific value of the corresponding grain pellet reward and have been found to elicit similar rates of Pavlovian and instrumental responding as a pellet reward in other experiments conducted in this lab. In all experiments involving liquids, the magazine was scrubbed with warm water and thoroughly dried between sessions to remove residual traces of the liquid reinforcer. To reduce neophobia to the reinforcers, one day prior to magazine training sessions all animals were pre-exposed to the reinforcers (10 g of pellets per animal and 25 ml of liquid reinforcer per animal) in their home cage.

### Magazine training

All animals received one session of magazine training for each experimental reinforcer with the following parameters: reward delivery was on an RT60 s schedule for 16 rewards. When necessary, sessions were separated by at least 2 hours and the order of reinforcer identity was counterbalanced between groups.

### Behaviour

CS responding was operationalized as the number of magazine entries during the CS period. PreCS responding was operationalized as the frequency of responding during the immediately preceding the CS period and was used as a measure of baseline responding to the testing context. PreCS responding was analysed separately, and any group differences identified and reported. Data were presented as CS – PreCS difference scores, which reflect discriminative responding to the CS. All data were analysed with mixed ANOVAs using R statistical software (Lenth et al., 2020; R Core Team (2020), 2020; Singmann et al., 2020), and significant interactions of interest were followed up with ANOVAs on the relevant subset of data, and simple effects with a Tukey family-wise error rate correction. Where relevant, planned linear and quadratic orthogonal trend contrasts and their interactions between groups were analysed to assess differences in rates of responding.

## Experiment 1: Pre-training OFC lesions

### Subjects

Subjects were forty-eight (N = 48) rats, tested in two cohorts. Cohort 1, n = 16 rats weighing between 280-361 g (M = 312.2 g) and cohort 2, n = 32 rats weighing between 271-328 g (M = 296.3 g).

### Training

#### Pavlovian Acquisition

Following magazine training, all rats received 21 sessions of Pavlovian acquisition training. Each session consisted of 16 presentations of a single auditory CS (a 15 s train of clicks) presented on a VT90s schedule (ranging from 60 to 120 s). A single pellet (US) was delivered at the termination of each CS. The session duration was 28 mins and animals were left in the chamber for an additional 2 mins before being removed. Animals received either one session per day, or two sessions per day separated by at least 2 hours.

#### Subgroup 1: General Satiety Pre-Feeding

At the end of acquisition training on day 21, a subgroup of animals (sham n = 8, lesion n = 8) were taken off food restriction and given 24 hours free access to their home cage food before further acquisition training on day 22. This session was rewarded as per acquisition training. At the end of day 22 animals were put back on food restriction and continued acquisition training.

#### Subgroup 2: Devaluation

Following initial Pavlovian acquisition of a single CS-US association, a subgroup of animals (sham n = 8, lesion n = 8) were re-trained with two novel unique CS-US associations intended to test devaluation in a taste aversion procedure.

#### Novel Acquisition

Novel acquisition of two unique CS-US associations was conducted with identical parameters to initial acquisition training, 2 session per day for 14 days, each session consisting of 16 trials consisting of a 15s CS co-terminating with reward with a vITI90s. Unlike initial acquisition the two CSs were an 80dB white noise and a 2800 Hz, 80 dB tone followed by either a single pellet or 20% w/v maltodextrin liquid (CS-US identities counterbalanced between animals).

#### Taste Aversion

Taste aversion took place in the devaluation chambers and involved 30 mins exposure to one US every day, alternating each day for 4 days. Following fee access to a US, animals were immediately injected i.p. with either 0.15M LiCl or 0.9% saline (15 mL/Kg). The outcome paired with nausea induced by injection of LiCl was designated the devalued outcome and the outcome paired with neutral saline injections was designated the non-devalued outcome (counterbalanced between animals). Following the final day of injections all animals were given a day of rest in their home cage to allow hunger levels to return to normal after taste aversion training.

#### Devaluation Test

Animals were tested with a single session of CS training except that no rewards were delivered i.e. in extinction. The magazine frequency measure that was available was not as sensitive to devaluation as a measure of duration, so only data from the first trial was analysed at test.

#### Locomotor Activity

At the end of the experimental procedures, all animals were assessed for locomotor activity over a 1-hour period.

### Histology and Group Allocation

Lesion damage is depicted in Figure 1-figure supplement 1. Lesion extent was judged by a trained observer blind to group allocation. A lesion was retained if there was evidence of significant bilateral damage constrained to LO or DLO. Animals were excluded if there was only unilateral LO/DLO damage, evidence of damage to the dorsal part of the anterior olfactory nucleus ventral to LO/DLO or if there was extensive damage to the white matter of the forceps minor of the corpus callosum. One lesioned animal did not recover from surgery, four lesion animals had only unilateral OFC damage, and one lesioned animal had extensive white matter damage. Forty-two animals were retained (N = 42, sham n = 24, lesion n = 18), of which subgroup 1 contained fifteen (*N* = 15; sham *n* = 8, lesion *n* = 7) and subgroup 2 contained thirteen (*N* = 13; sham *n* = 8, lesion *n* = 5).

### PreCS Analysis

Analysis of the PreCS period using a Group (sham, lesion) x Block (1-7) mixed ANOVA revealed that responding was significantly higher in the lesion group than the sham group (main effect of Group *F*_(1, 40)_ = 7.24, *p* = .01). Furthermore, while responding increased over blocks

(main effect of Block *F*_(6, 240)_ = 20.37, *p* < .001; positive linear trend *F*_(1, 40)_ = 33.18, *p* < .001), this increase was greater in the lesion than the sham group (Block x Group interaction *F*_(6, 240)_ = 2.52, *p* = .02; linear trend interaction *F*_(1, 40)_ = 5.34, *p* = .03). During the first block PreCS responding was similar between groups (Sham M = 2.07, SD = 0.60; Lesion M = 2.13, SD = 0.90), by the final block PreCS responding was higher in the Lesion group (M = 4.30, SD = 1.95) than the sham group (M = 2.76, SD = 2.30).

## Experiment 2: Post-training muscimol inactivation

### Subjects

Subjects were thirty-two (total N = 32) male Long Evans rats (Monash Animal Services, Gippsland, Victoria, Australia) approximately 4 months old, weighing between 285-350 g (M = 319.7 g).

### Pavlovian Acquisition

Animals were given 9 sessions, 1 session per day, of Pavlovian acquisition training with session parameters identical to those described in Experiment 1. This number of sessions was chosen because the effect of pre-training lesions appeared after around 9 session in Experiment 1. Briefly, each session consisted of a VT90s ITI with 16 trials consisting of a 15s click CS co-terminating with a single pellet US. Following the final day of training all animals were taken off food restriction and received surgical implantation of guide cannulae.

### Post-Training

#### Pre-Infusion

Following post-operative recovery animals were returned to food restriction for a day before receiving a further 2 days of acquisition training as per pre-training. However, immediately prior to entering the chamber all animals received a dummy infusion.

#### Infusion

Animals were assigned to one of two infusion groups such that performance there were no differences between groups on the final day of pre-infusion acquisition. For the next 4 days, all animals received an infusion of saline or Muscimol immediately prior to entering the testing chamber for a Pavlovian acquisition session.

#### Post-Infusion

On the final 2 days of training all animals received a further 2 days of acquisition training immediately preceded by a dummy infusion.

### Histology and Group Allocation

Cannulae placements are illustrated in Figure 2-figure supplement 2. One animal did not recover from surgery and was excluded. Three animals were excluded because of the cannulae assembly detaching from the skull. A further 3 animals were excluded because of failing to consume the pellets after recovery from surgery. One animal from the muscimol group was excluded from analysis because of a cannula tip embedded within the white matter of the forceps minor of the corpus callosum. Therefore, a total of 8 animals were excluded leaving *N* = 24 (saline *n* = 12, muscimol *n* = 12).

### PreCS Rates

PreCS baseline responding did not differ between infusion groups across training and justified the use of CS-preCS difference scores for analyses of discriminative responding. In particular, during the infusion period a Group x Day (4 days) mixed ANOVA on preCS responses revealed a significant effect of Day (*F*_(3, 66)_ = 5.95, *p* = .001) but no significant effect of Group (*F*_(1, 22)_ = 0.01, *p* = .93) or Group x Day interaction (*F*_(3, 66)_ = 0.41, *p* = .741). PreCS response rates on these days were, saline *M* = 0.70, *SD* = .48, muscimol *M* = 0.72, *SD* = .48.

## Supplementary Experiment 1: Post-Training OFC lesions

### Methods

#### Subjects

Subjects were twenty-four (total N = 24) male Long Evans rats (Monash Animal Services, Gippsland, Victoria, Australia) approximately 4 months old, weighing between 317-369 g (M = 338.9 g).

#### Pre-lesion Training

##### Pavlovian Acquisition

All animals received 9 days of Pavlovian acquisition training, 1 session per day. On the final day of training all animals were removed from food restriction for at least 24 hours before receiving sham or excitotoxic lesions of the OFC. Lesion conditions were pseudo-randomly assigned to animals such that group performance was matched on the final day of acquisition and an equal number of animals were assigned to each lesion condition in each homecage.

#### Post-lesion Training

##### Pavlovian Acquisition

Following post-operative recovery all animals were returned to food restriction for 24 hrs before receiving an additional 9 days of acquisition training.

#### Histology and Group Allocation

Lesion damage is depicted in Figure 2-figure supplement 3. Lesion extent was judged by a trained observer blind to group allocation. A lesion was retained if there was evidence of significant bilateral damage constrained to LO or DLO. Animals were excluded if there was only unilateral LO/DLO damage, evidence of damage to the dorsal part of the anterior olfactory nucleus ventral to LO/DLO or if there was extensive damage to the white matter of the forceps minor of the corpus callosum. Three lesion animals had only unilateral OFC damage and were excluded from analysis (final *N* = 21; sham *n* = 12, lesion *n* = 9).

#### PreCS Responding

PreCS levels of responding did not differ between groups across days of training, and on the final block of 3 days (post-operative) response rates (15s) were sham *M* = 2.55, *SD* = 2.03, lesion *M* = 2.74, *SD* = 0.94. A mixed Group x DayBlock (6 blocks of 3 days) ANOVA on preCS responding supported this observation with only a significant main effect of DayBlock (*F*_(5, 95)_ = 11.52, *p* < .001, effect of Group and Group x DayBlock interaction *F* < 1.00, *p* > .81).

## Supplementary Experiment 2: OFC inactivation early in acquisition

### Subjects

Subjects were thirty-two (total N = 16) male Long Evans rats (Monash Animal Services, Gippsland, Victoria, Australia) approximately 4 months old, weighing between 321-399 g (M = 357.4 g).

### Surgery

Surgical implantation of cannulae occurred prior to any behavioural training.

#### Pavlovian Acquisition

Animals were given 10 sessions, 1 session per day. Briefly, each session consisted of a VI 200s ITI with 16 trials consisting of a 10s light CS (illumination of the house light at the back of the chamber) co-terminating with a single pellet US. Subjects received mock infusions on days 3 and 4, and either Saline or Muscimol was infused prior to entering the chamber on days 5-9. On day 10 all animals received a mock infusion.

### Histology and exclusions

Cannulae placements are illustrated in Figure 3-figure supplement 1. One rat in the Muscimol condition had a blocked guide cannulae and was excluded from experimental analysis. Final numbers N = 15 (Muscimol n = 7, Saline n = 8).

### PreCS Rates

PreCS responding did not differ between infusion groups across the 10 days of Pavlovian conditioning (Group *F*_1,13_ = 2.72, *p* = .12; Day *F*_9,117_ = 1.49, *p* = .16; Group x Day *F*_9,117_ = 2.72, *p* = .25).

## Experiment 3: OFC inactivation prior to associative blocking

### Subjects

Subjects were thirty-two (total N = 32) male Long Evans rats (Monash Animal Services, Gippsland, Victoria, Australia) approximately 4 months old, weighing between 299-395 g (M = 331.5 g).

### Surgery

Surgical implantation of cannulae occurred prior to any behavioural training.

### Training

The design of the experiment was such that 4 CSs were designated as cues A, B, C and D. Cues A and C were always visual cues, either darkness caused by extinguishing the houselight or flashing panel lights (5Hz; Figure 3A). Cues B and D were always auditory cues, either an 80dB white noise or a 5Hz train of clicks. Throughout all training sessions the house light was always illuminated unless it was extinguished to act as a visual cue. All cues lasted 10s and co-terminated with the delivery of the US, 2 pellets delivered consecutively 0.25s apart. The identity of the cues was counterbalanced between subjects except that A and C were always visual cues and B and D were always auditory cues. Simultaneous audio-visual compounds were designated as AB and CD. Pavlovian training sessions were always 56 mins long such that there were 16 trials with a vITI 200s (range 100 to 300s); animals were left in the chambers for an additional 2 mins before being removed.

#### Food Restriction and Magazine Training

Magazine training sessions consisted of an RT120s reward delivery schedule for 16 rewards. Each reward consisted of 2 pellets delivered to the magazine 0.25s apart.

#### Stage 1

Stage 1 acquisition involved 10 days of acquisition to cue A, 16 trials per session. On days 1-4 of training all animals received dummy infusions to familiarise them to the infusion procedure. Animals were then split into two groups with matched performance on day 4. On days 5-10 all animals received an infusion of saline or muscimol immediately prior to entering the test chambers.

#### Pre-exposure

On day 11 all rats received pre-exposure to auditory cues B and D, 4 non-rewarded presentations of each cue vITI 200s. This was done to minimise novelty to the auditory cues during compound training in stage 2. All animals received dummy infusions prior to the session.

#### Stage 2

On days 12-14 all animals received stage 2 audio-visual compound training. Sessions involved 8 presentations of compound AB and 8 presentations of CD (pseudo randomly presented such that a compound was never repeated more than 2 times in a row). The compounds were rewarded with 2 pellets, the same US that was used in stage 1. All animals received dummy infusions prior to each session.

#### Test

On day 15 and 16 all animals were tested in extinction for responding to the target auditory cue B and the overshadowing control cue D (8 presentations of each cue, pseudorandom trial order, vITI 200s). All animals received dummy infusions prior to each session.

#### Re-acquisition

On days 17-19, all animals received re-acquisition training to cue B (16 trials per session) to test for differences in rates of re-acquisition to the blocked cue. On days 20-21 animals were tested for re-acquisition to cue A (16 trials per session) to test for differences in the rate of re-acquisition to the blocking cue.

## Results

### Histology and Group Allocation

Cannulae placements are illustrated in Figure 3-figure supplement 3. 1 animal failed to consume pellets throughout the experiment and was excluded from testing. One animal from the muscimol group lost its cannula assembly during the infusion period and was excluded from testing. One animal in the muscimol group was euthanized due to severe illness. A further 2 animals were excluded after histological analysis revealed that the cannulae were only unilaterally targeting DLO and LO. Therefore, a total of 6 animals were excluded leaving *N* = 26 (saline *n* = 13, muscimol *n* = 13).

### PreCS Responding

Baseline levels of responding did not differ between groups during training, and on the final day of infusions (day 10 of stage 1) preCS response rates (10s) were saline *M* = 0.122, *SD* = 0.24, muscimol *M* = 0.67, *SD* = 0.87. These observations were supported by mixed Group x Day ANOVAs on preCS responding in stage1 suggesting that there were no group differences on days 1-4 prior to infusion (all *F* < 1.69, *p* > .21) or on days 5-10 during infusions (significant main effect of Day *F*_(5, 120)_ = 15.21, *p* < .001, all remaining *F* < 1.00, *p* > .50).

## Supplementary Figures

**Figure 1-figure supplement 1.**
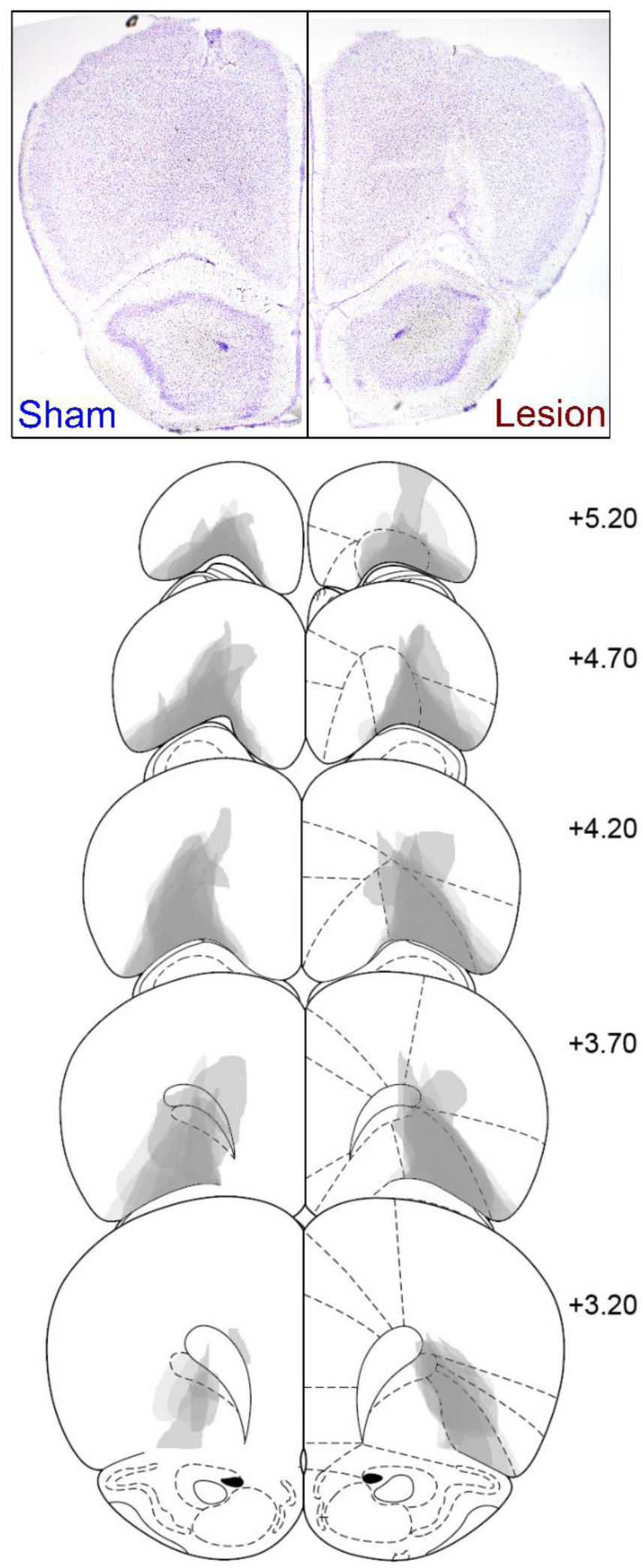
Photomicrograph of representative OFC lesion damage (top) in the sham (left) and lesion (right) groups. Coronal slice located approximately +4.20 mm anterior to bregma. Semi-transparent grey patches (bottom) represent lesion damage in each subject, and darker areas represent overlapping damage across multiple subjects. Coronal sections are identified in mm relative to bregma (Paxinos and Watson, 1997).

**Figure 1-figure supplement 2.**
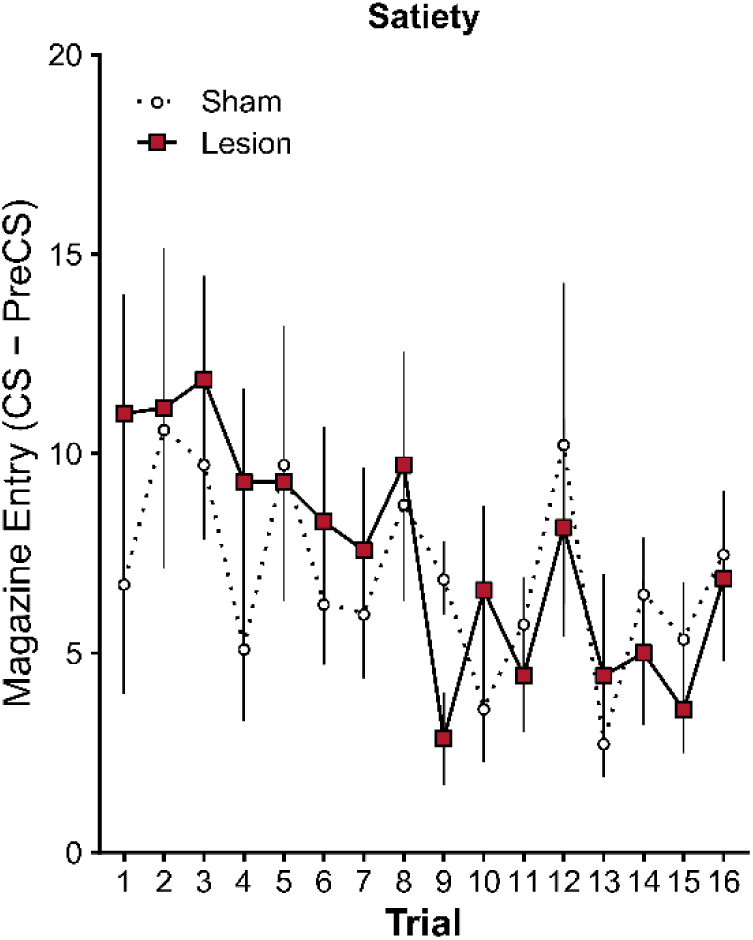
Responding on each trial within-session is presented following a general satiety manipulation (session average presented in Figure 1B). Conditioned responding was not significantly different between the lesion and sham group, and this is evident on trial 1 prior to contact with the reward (*t*(13) = 1.04, *p* = .317). Error bars depict ± SEM.

**Figure 1-figure supplement 3.**
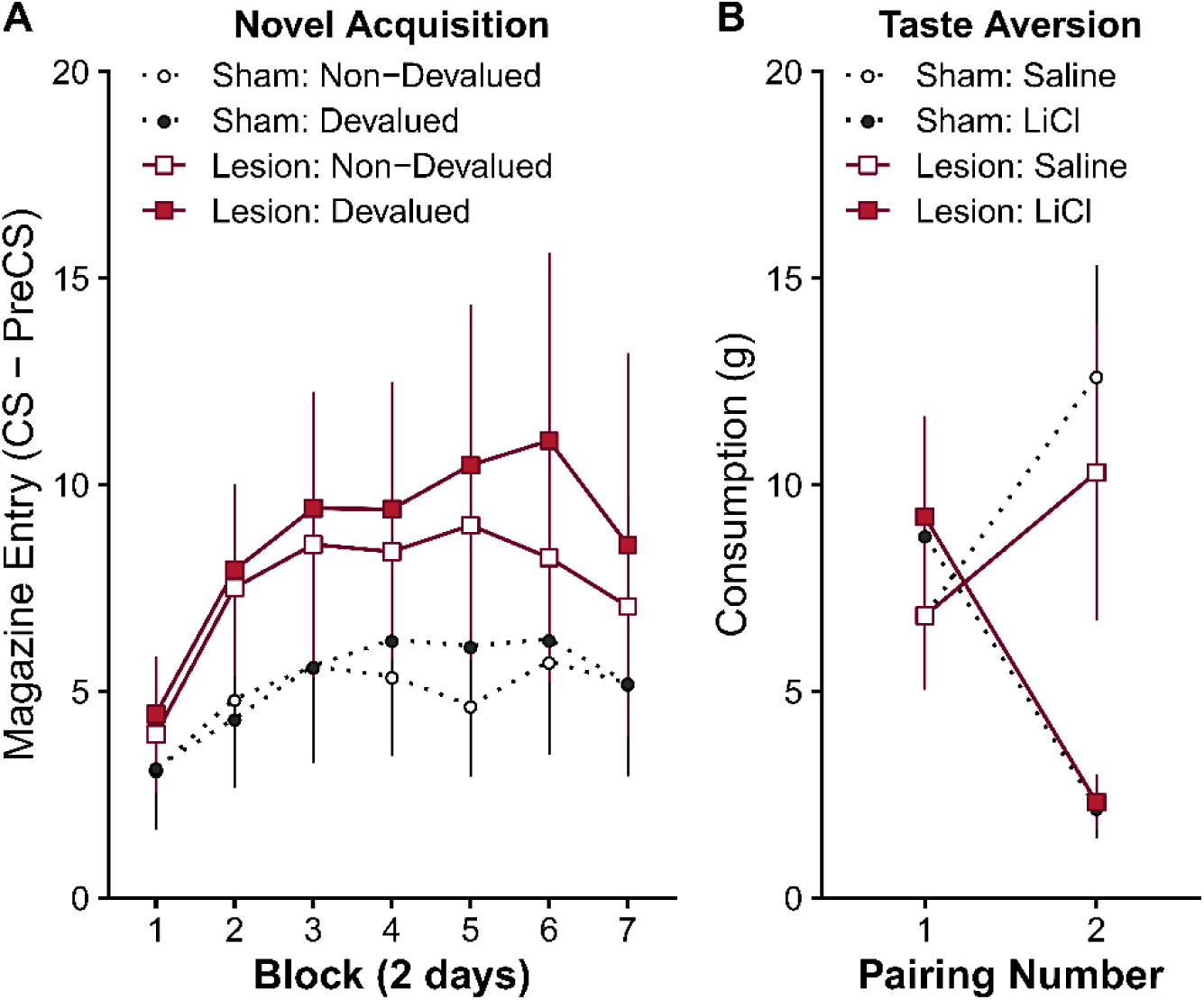
(**A**) CS-PreCS response levels during the acquisition of two novel and distinct cue-outcome pairings. One outcome was designated for subsequent devaluation, and responding is presented separately for the cue that predicts the to-be devalued and non-devalued outcomes.While mean response rates suggested elevated responding in the lesion group, this observation was not supported statistically (main effect of Block *F*(6,66) = 3.96, *p* = .002, all remaining effects *F* < 0.98, *p* > .343). It is likely that the elevated responding in the lesion group did not reach significance due to the reduced number of subjects in the lesion group. (**B**) Average consumption (g) of the outcome followed immediately followed by an injection of saline or LiCl to establish a selective taste aversion. All animals acquired a significant taste aversion to the outcome paired with post-consumption LiCl injections (significant Injection x Pairing interaction *F*(1,11) = 49.58, *p* < .001, all remaining effects *F* < 2.03, *p* > .182), which did not significantly differ between lesion groups. Specifically, consumption of the outcome paired with LiCl significantly decreased between pairing 1 and 2 (LiCl: pairing 1 vs 2 *t*(11) = 4.26, *p* = .001), whereas consumption of the outcome paired with saline increased (saline: pairing 1 vs 2 *t*(11) = −3.74, *p* = .003).Error bars depict ± SEM.

**Figure 2-figure supplement 1.**
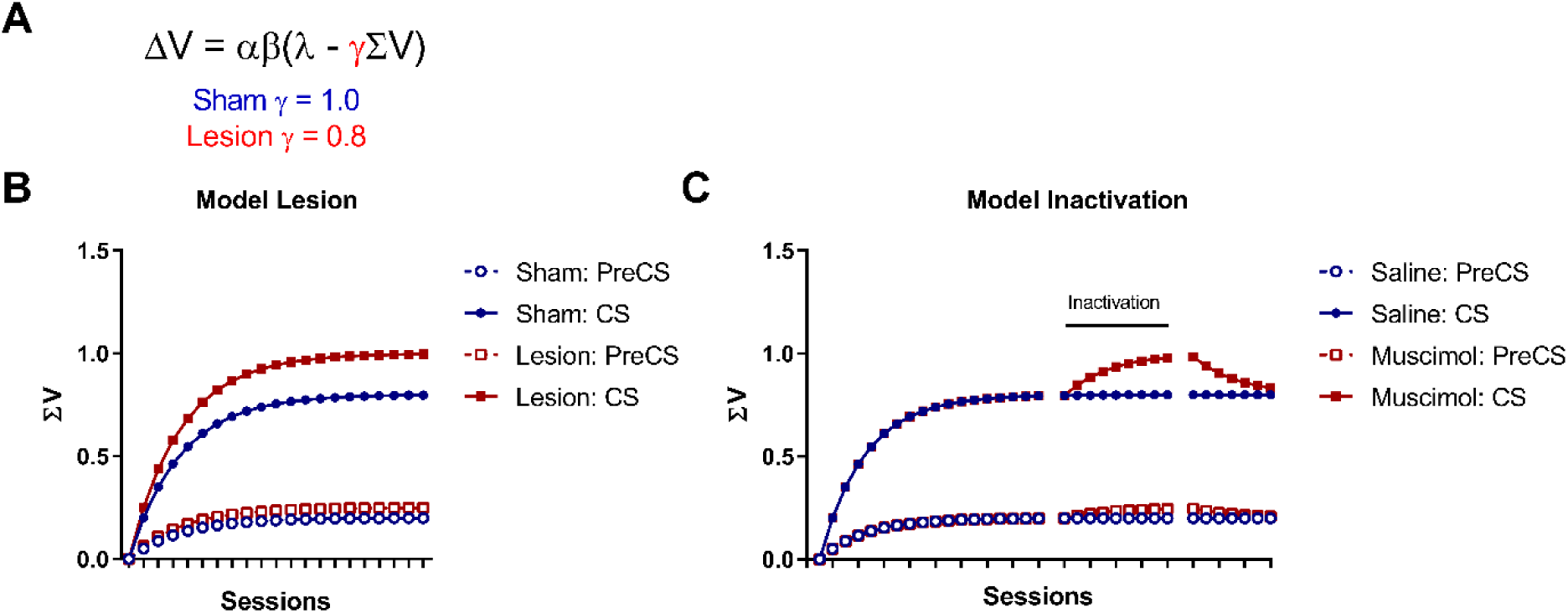
**Model predictions of LO lesions as a partial loss of outcome expectancy. (A)** The model predictions for simple Pavlovian acquisition using a standard Rescorla-Wagner learning rule and following modelled lesion deficits as a partial loss of outcome expectancy information specific to learning. Using the following update rule: Δ*V* = *αβ*(*λ* − γΣV), the change in associative strength (Δ*V*) was calculated on a trial by trial basis for the CS and the context (preCS) periods. Parameters used were *αβ* = .05 for the context and .2 for the CS, *λ* = 1, and γ = 0.8 for lesions and γ = 1.0 for the control condition. **(B)** Modelled performance following pre-training sham or LO lesions. **(C)** Modelled performance following temporary LO inactivation via muscimol after initial training. One potential account of the heightened responding observed during simple acquisition following LO lesions is enhanced cue-outcome learning. However, given the hypothesized role of the OFC in the representation of outcome expectancy value (Baxter, Parker, Lindner, Izquierdo, & Murray, 2000; Pears, Parkinson, Hopewell, Everitt, & Roberts, 2003; Schoenbaum, Roesch, Stalnaker, & Takahashi, 2009; Takahashi et al., 2009, 2011), it is not immediately clear how this could be the case. The Rescorla-Wagner model of the acquisition of associative value suggests that in the absence of a representation of the expected value of an outcome, no conditioned responses should be expressed. In fact, in most associative learning theories the rules that govern performance are some function of the expected value of the outcome (Esber & Haselgrove, 2011; LePelley, 2004; Mackintosh, 1975; Nasser, Calu, Schoenbaum, & Sharpe, 2017; Pearce & Hall, 1980; Rescorla & Wagner, 1972; Sutton & Barto, 1998). However, OFC damage may not abolish all outcome expectancy information but instead might degrade some aspect (e.g. sensory specific properties) of the outcome expectancy information made available at the time of calculating the prediction error (e.g. mid brain Dopamine neurons; Takahashi et al., 2011). This would assume that the expected value used in prediction error learning may not necessarily be the same as the full expected value used to govern performance. To model this mathematically, a constant (γ) is used to represent the proportion of the outcome expectancy information that was available for learning (<u>Figure 2-Figure supplement 1A</u>). The strength/value (V) of the learned association changes in the following manner:

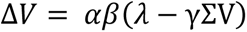

Such that Δ*V* is the change in associative strength on a given trial; *αβ* are learning rate parameters based on the properties of the cue and outcome respectively such that 0 ≤ *α* ≤ 1 and 0 ≤ *β* ≤ 1; *λ* is the experienced value of the outcome on a given trial; ΣV is the sum of the associative strength i.e. the expected value. The constant γ represents the proportion of ΣV available for learning such that 0 ≤ γ ≤ 1. In a healthy control animal 100% (γ = 1) of the available outcome expectancy information is available to guide learning. Consequently, further learning will stop once *λ* = ΣV as the current expected value of the outcome fully accounts for the actual value of the outcome. Following OFC lesions some of this information is lost and, for example, 80% (γ = 0.8) of the available outcome expectancy information is available to guide learning. Therefore, learning will continue beyond the point at which *λ* = ΣV and will only stop once *λ* = γΣV. Therefore, the asymptote of conditioned behaviour (expressed as some function of ΣV) will be determined by 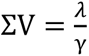, therefore asymptotic conditioned behaviour will be higher following a lesion in this model (γ < 1) than an intact OFC (γ = 1). The following simulation of this model (with the lesion parameter set at γ = 0.8) reveals a pattern of acquisition that is similar at the start of acquisition but diverges considerably towards the end of acquisition. This pattern strongly resembles the findings in Figure 1A with a range of values of γ > 0.6. This provides prima facie evidence of how well the model accounts for the present data; however, the most valid test of the model would be an experimental test of a hypothesis generated from the model. This model predicts that if acquisition initially occurs with a functional LO until asymptote (i.e. ΣV = *λ*) and no more learning is evident, inactivation of OFC should allow learning to temporarily increase above asymptote because *λ* < γΣV. Informally, the inactivation of LO will cause the outcome to no longer be fully predicted by the CS because part of the expectation is now missing. Similarly, if the function of LO is returned, then any new learning will extinguish until *λ* = ΣV again. This prediction is modelled in (**C**). It is noteworthy that modelling the effect of LO damage in this way produces identical predictions in a number of other associative learning models (Esber & Haselgrove, 2011; Mackintosh, 1975; Pearce & Hall, 1980; Rescorla & Wagner, 1972) and therefore may also affect other aspects of learning such as attention. Furthermore, even though the hypothesis was generated using the current mathematical model, the prediction and underlying mechanism would be the same for any interpretation of the increased responding in Figure 1A as increased cue-outcome learning. If OFC damage increases learning, then transient inactivation of OFC should temporarily allow learning to increase.

**Figure 2-figure supplement 2.**
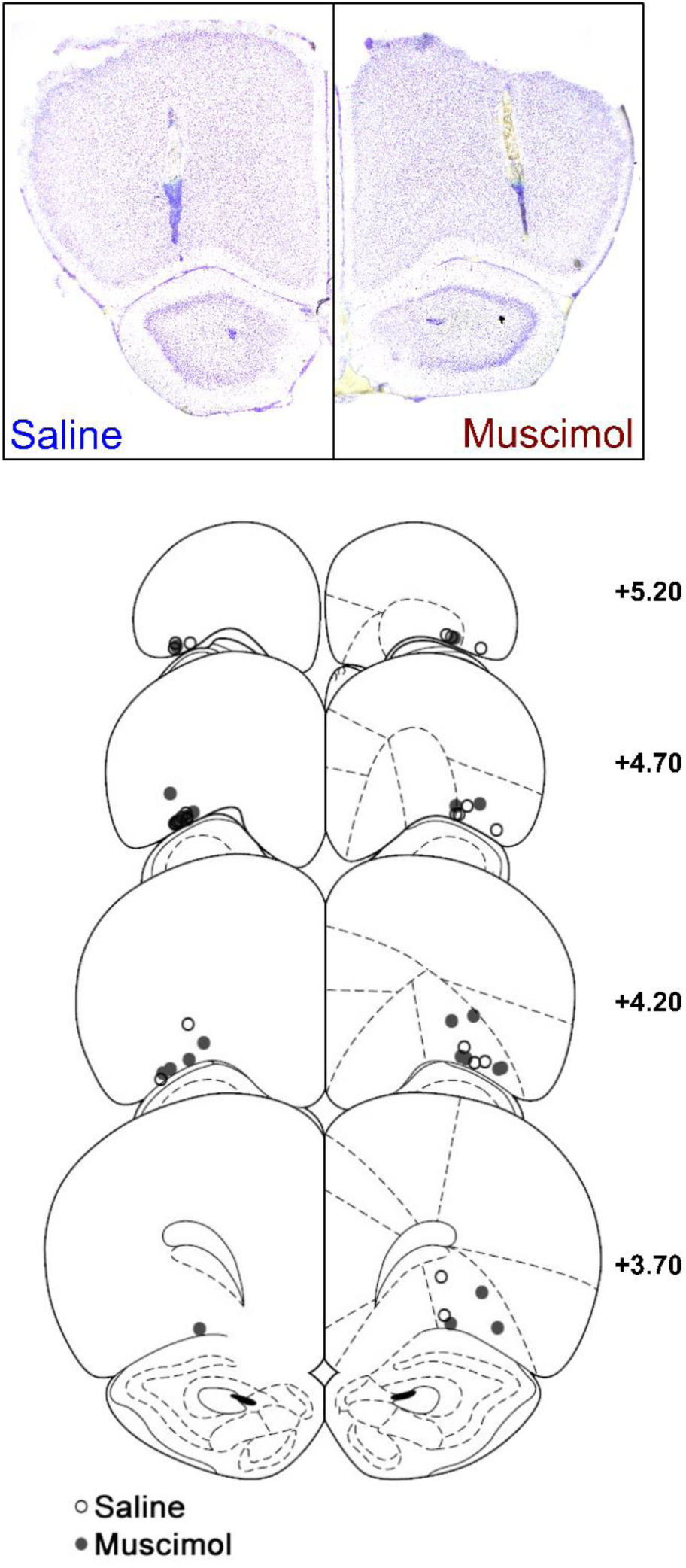
Photomicrograph of representative OFC cannulae placement (top) in the saline (left) and muscimol (right) infusion groups. Coronal slice located approximately +4.20 mm anterior to bregma. Cannulae tip (bottom) location of all subjects in the saline (empty circles) and muscimol (filled circles) groups. Coronal sections are identified in mm relative to bregma (Paxinos and Watson, 1997).

**Figure 2-figure supplement 3.**
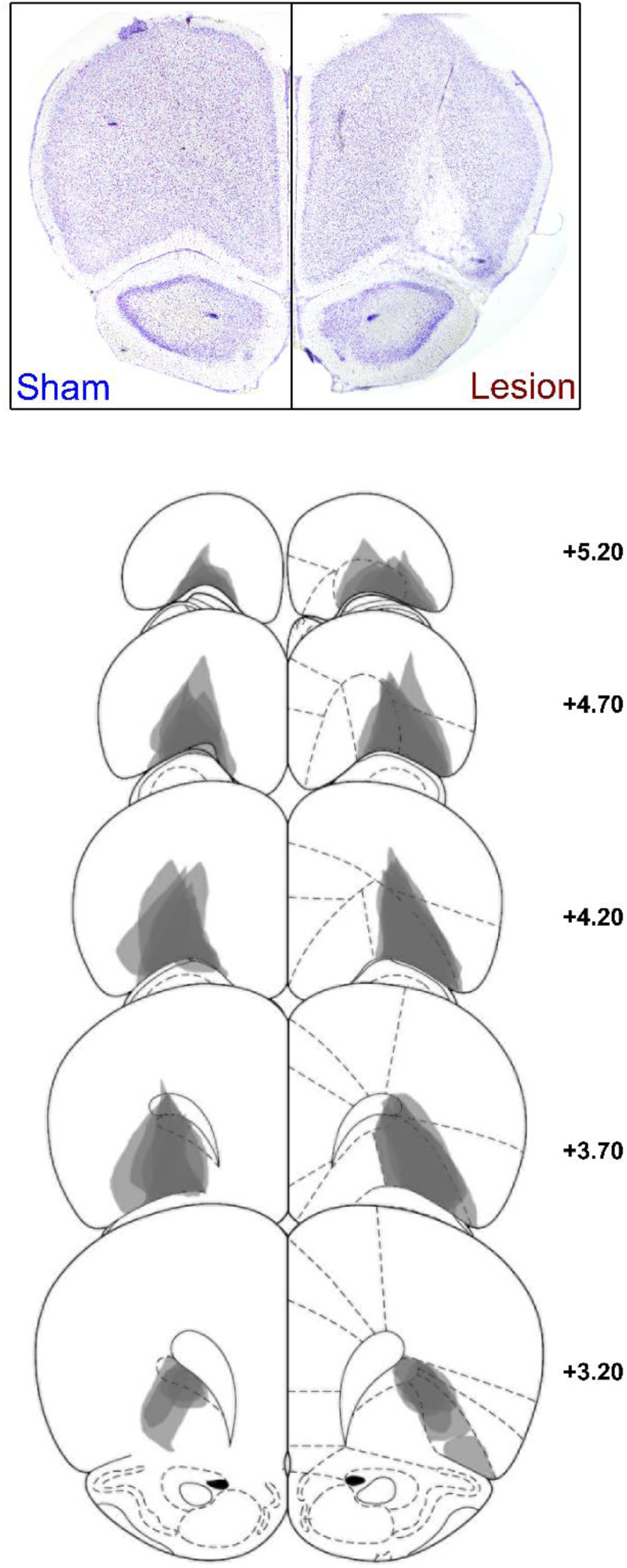
Supplementary Experiment 1: Photomicrograph of representative OFC lesion damage (top) in the sham (left) and lesion (right) groups following post-training lesions (behavioural data depicted in <u>Figure 2-figure supplement 4)</u>. Coronal slice located approximately +4.20 mm anterior to bregma. Semi-transparent grey patches (bottom) represent lesion damage in each subject, and darker areas represent overlapping damage across multiple subjects. Coronal sections are identified in mm relative to bregma (Paxinos and Watson, 1997).

**Figure 2-figure supplement 4.**
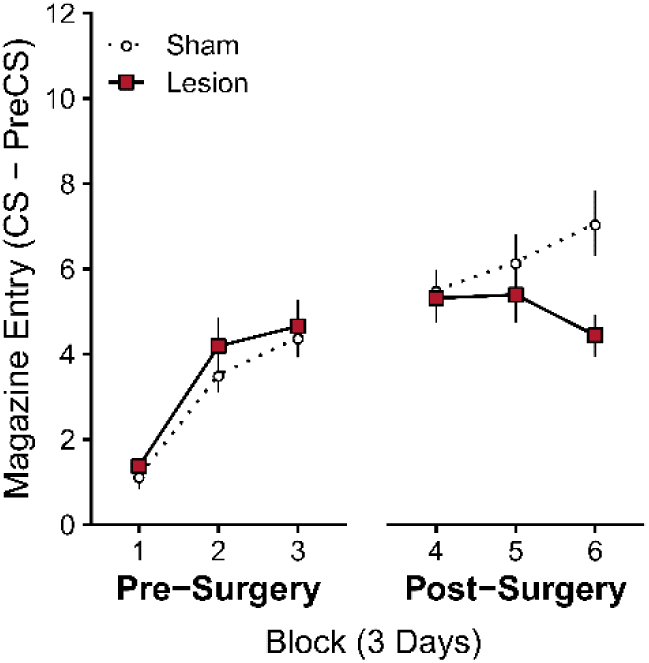
Supplementary Experiment 1: Post-training OFC lesions suppress Pavlovian acquisition behaviour. Rates of discriminative responding (CS-PreCS) during initial acquisition (Pre-Surgery, Blocks 1-3), and following OFC or sham lesions (Post-Surgery, Blocks 4-6). Data summarized in blocks of 3 days. Histological characterization of lesion extent depicted in <u>Figure 2-figure supplement 3</u>. Error bars depict ± SEM. We trained a new cohort of animals on this simple Pavlovian cue-outcome task for 9 days, and then performed post-training excitotoxic or sham OFC lesions before continuing with acquisition (lesion extent depicted in Figure 2-figure supplement 3). Prior to surgery, animals acquired responding to the cue (Pre-Surgery; significant main effect of Block *F*(2,38) = 61.98, *p* < .001, but no main effect of Group *F*(1,19) = 0.49, *p* = .492, or Group x Block interaction *F*(2,38) = 0.31, *p* = .738). After surgery, the sham group continued to acquire responding, but the lesion group did not (Post-Surgery; significant Group x Block interaction *F*(2,38) = 6.25, *p* = .005 , but no main effect of Group *F*(1,19) = 2.21, *p* = .154, or Day *F*(2,38) = 0.66, *p* = .525). Responding in the sham control group was significantly higher than the lesion group in the final block of 3 days (Block 4 *t*(19) = −0.25, *p* = .806, Block 5 *t*(19) = −0.80, *p* = .434, Block 6 *t*(19) = −2.65, *p* = .016). Furthermore, further acquisition post-surgery was completely abolished in the lesion group (Lesion: no linear trend over Blocks 4-6 *t*(19) = −1.42, *p* = .172), but continued in the sham control group (Sham: significant positive linear trend over Blocks 4-6 *t*(19) = 2.93, *p* = .009. Therefore, both post-training lesions and inactivation of OFC function disrupted Pavlovian acquisition. To facilitate comparisons between experiments, CS-PreCS response rates on Block 6 in the present experiment were Sham: M = 9.61, SD = 3.88, Lesion: M = 7.18, SD = 1.74. The terminal levels of responding in the sham group are similar to those of the saline group in Figure 2, and the sham group in Figure 1A which used identical session parameters. This suggests that the present findings are also unlikely to be due to abnormally elevated levels of responding in the control groups in any one of these experiments.

**Figure 3-figure supplement 1.**
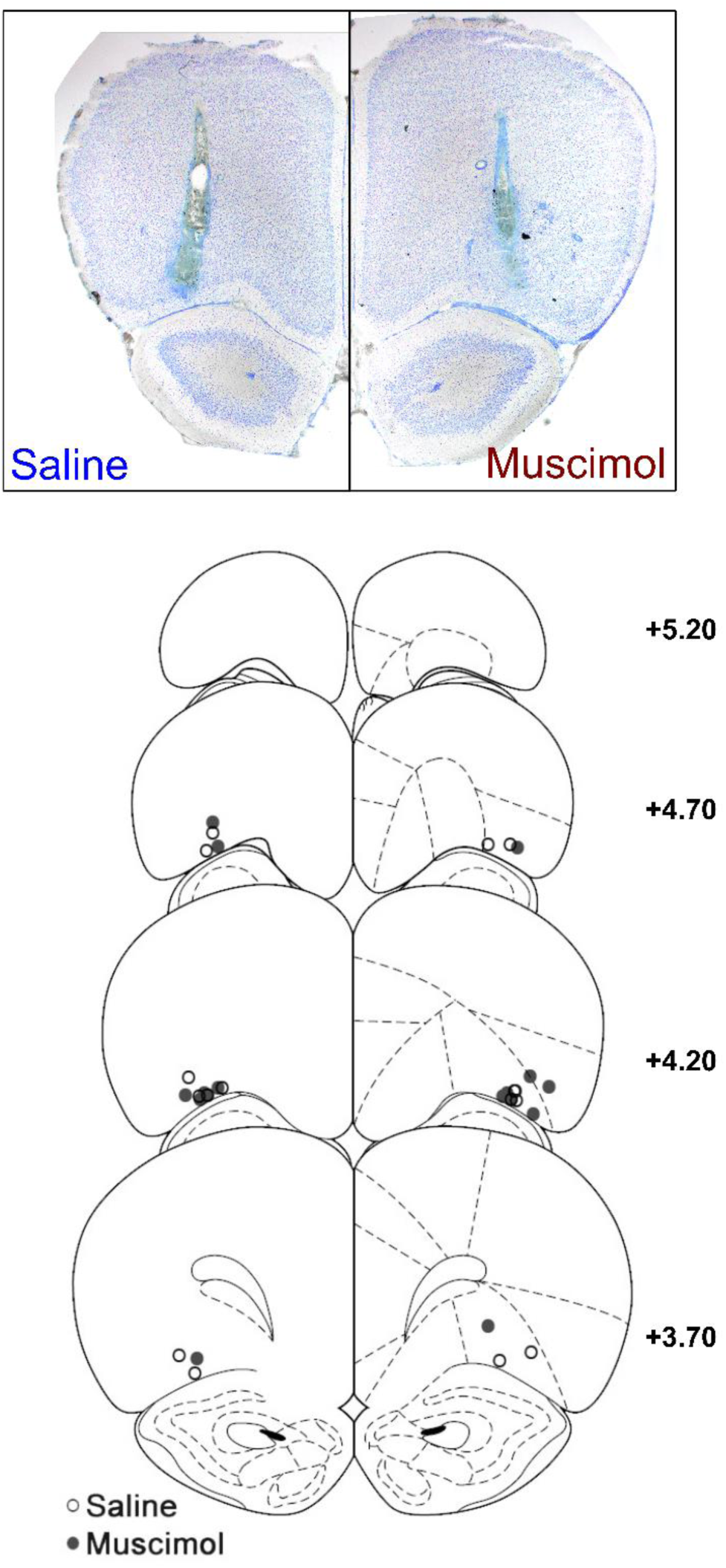
Supplementary Experiment 2: Photomicrograph of representative OFC cannulae placement (top) in the saline (left) and muscimol (right) infusion groups (associated behavioural data presented in <u>Figure 3-figure supplement 1</u>). Coronal slice located approximately +4.20 mm anterior to bregma. Cannulae tip (bottom) location of all subjects in the saline (empty circles) and muscimol (filled circles) groups. Coronal sections are identified in mm relative to bregma (Paxinos and Watson, 1997).

**Figure 3-figure supplement 2.**
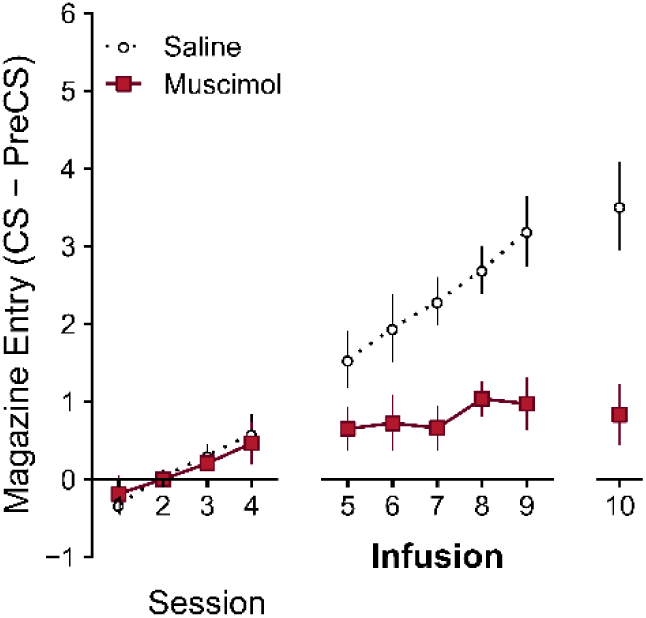
Supplementary Experiment 2: Post-training OFC inactivation early in training suppresses Pavlovian acquisition behaviour, and this suppression persists in the absence of OFC inactivation. Rates of discriminative responding (CS-PreCS) during initial acquisition (sessions 1-4), following intra-OFC infusion of muscimol or saline (sessions 5-9), and without infusion (session 10). Cannulae placements depicted in <u>Figure 3-figure supplement 1</u>. Note that the task presented in Figure 1 and Figure 2 employed a 15s auditory CS, whereas a 10s visual CS was used in the present figure and during the associative blocking experiment presented in Figure 3. Stimulus modality, salience, and duration differences account for the different levels of conditioned responding between these experiments. Error bars depict ± SEM. Prior to drug infusions, all animals acquired responding to the cue (Figure 2C, Days 1-4; Significant main effect of Day *F*(3,39) = 9.42, *p* < .001, but no main effect of Group , or Group x Day interaction *F*(3,39) = 0.30, *p* = .826). However, OFC inactivation during the next 5 days of conditioning significantly impaired acquisition in the muscimol group (Days 5-9; Significant main effect of Group *F*(1,13) = 12.57, *p* = .004, Day *F*(4,52) = 7.72, *p* < .001, and Group x Day interaction *F*(4,52) = 3.05, *p* = .025). Responding in the muscimol group was significantly lower than the saline group on days 7-9 (Muscimol vs Saline: Day 5 *t*(13) = −1.85, *p* = .087, Day 6 *t*(13) = −2.10, *p* = .056, Day 7 *t*(13) = −3.77, *p* = .002, Day 8 *t*(13) = −4.17, *p* = .001, Day 9 *t*(13) = −3.80, *p* = .002). Again, this deficit was characterised by significant acquisition over days in the saline group that was abolished in the muscimol group (positive linear trend over days 5-9; Saline *t*(13) = 6.59, *p* < .001, Muscimol *t*(13) = 1.45, *p* = .171). Finally, this reduction in responding persisted on day 10 when all rats were tested without infusion (Day 10; *t*(12.14) = 3.88, *p* = .002). In contrast to OFC inactivation later in acquisition (Figure 2), disrupting OFC activity early in learning suppressed performance which persisted when the OFC was active again. These findings suggest that OFC inactivation early in training disrupted acquisition learning rather than just behavioural performance.

**Figure 3-figure supplement 3.**
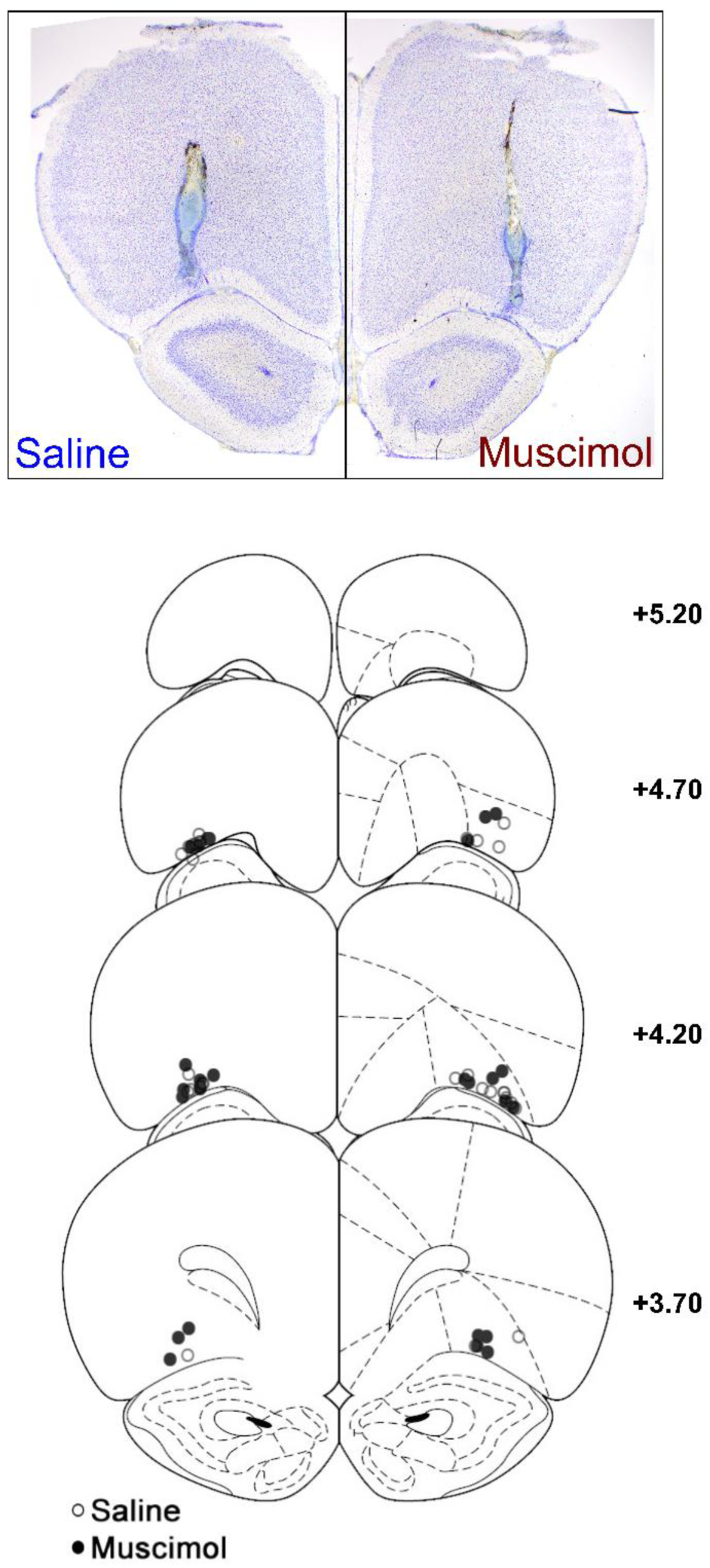
Photomicrograph of representative OFC cannulae placement (top) in the saline (left) and muscimol (right) infusion groups (associated behavioural data in Figure 3). Coronal slice located approximately +4.20 mm anterior to bregma. Cannulae tip (bottom) location of all subjects in the saline (empty circles) and muscimol (filled circles) groups. Coronal sections are identified in mm relative to bregma (Paxinos and Watson, 1997).

**Figure 3-figure supplement 4.**
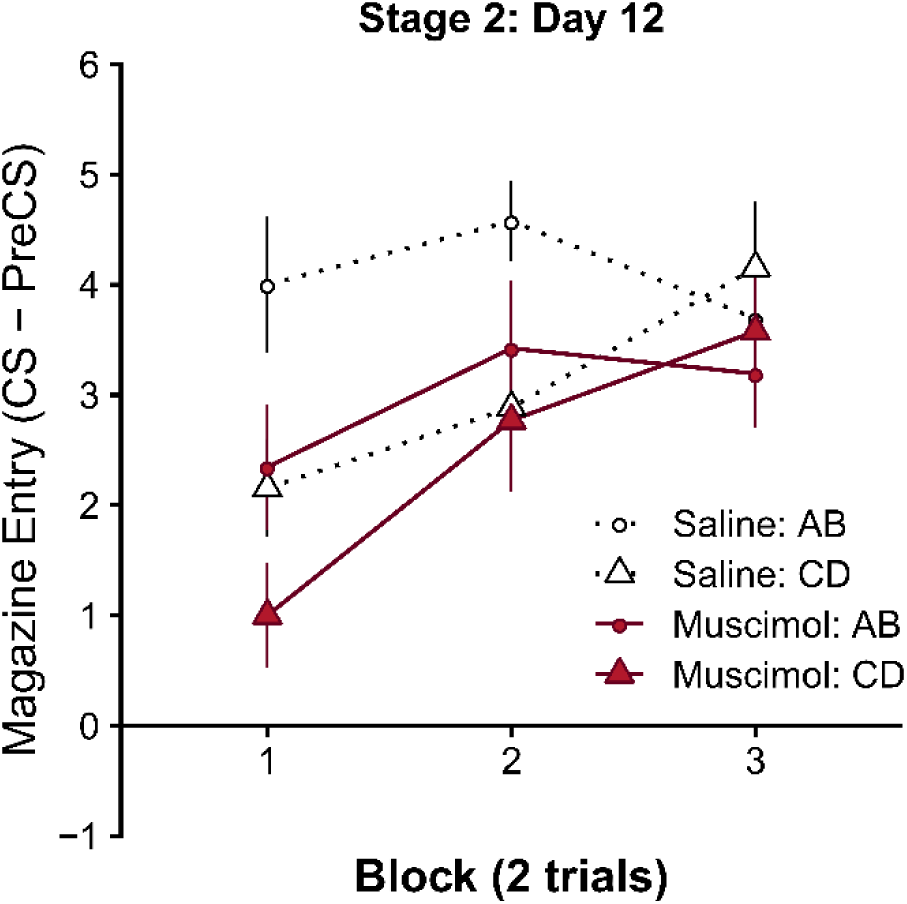
Within-session responding on Day 12, the first session of Stage 2 blocking (session average shown in Figure 3C). Rates of discriminative responding (CS-PreCS) presented in blocks of 2 trials. The effect of muscimol inactivation in Stage 1 is evident at the start of the session i.e. a significant reduction in responding relative to the saline group. First 2 trials, significant main effect of Group *F*(1,24) = 8.67, *p* = .007, and Cue *F*(1,24) = 7.61, *p* = .011, but no Group x Cue interaction *F*(1,24) = 0.19, *p* = .670). Error bars depict ± SEM.

**Figure 3-figure supplement 5.**
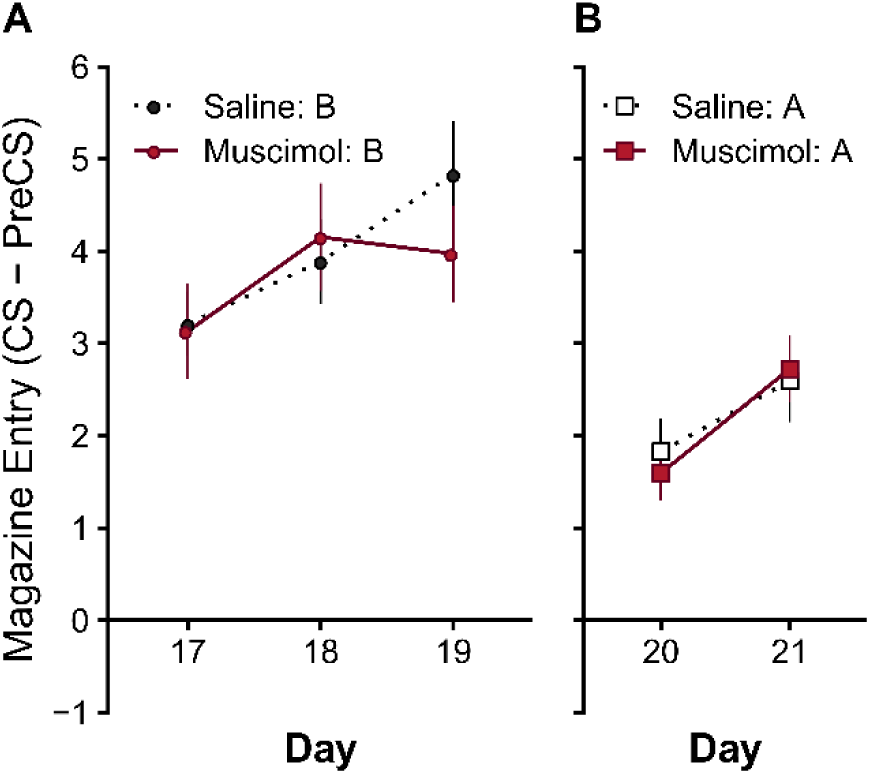
Reacquisition to cues A and B provide a test of whether the successful blocking effect observed in the saline and muscimol groups was the result of different underlying attentional strategies. Down-regulation of attention to a cue can result in retardation of subsequent acquisition (Mackintosh, 1975; Papini & Bitterman, 1993; Pearce & Hall, 1980). There were no differences in the rates of learning to the blocked cue B or to the blocking cue A. (**A**) The rate of re-acquisition to cue B (day 17-19) and (**B**) cue A (day 20-21) did not differ between groups (Cue B: Significant main effect of Day *F*(2,44) = 9.54, *p* < .001, but no main effect of Group *F*(1,22) = 0.12, *p* = .727, or Group x Day interaction *F*(2,44) = 1.99, *p* = .148. Cue A:Significant main effect of Day *F*(1,22) = 54.73, *p* < .001, but no main effect of Group *F*(1,22) = 0.01, *p* = .919, or Group x Day interaction *F*(1,22) = 2.02, *p* = .169). Note: Due to experimenter error one animal in each group was tested with the wrong counterbalancing and excluded from the analysis of re-acquisition data (remaining N = 24, saline n = 12, muscimol n = 12). Error bars depict ± SEM.

## References

1. Balleine, B. W., Leung, B. K., & Ostlund, S. B. (2011). The orbitofrontal cortex, predicted value, and choice. Ann N Y Acad Sci, 1239, 43–50. https://doi.org/10.1111/j.1749-6632.2011.06270.x

2. Barreiros, I. V, Ishii, H., Walton, M. E., & Panayi, M. C. (2021). Defining an orbitofrontal compass: functional and anatomical heterogeneity across anterior-posterior and medial-lateral axes. Behavioral Neuroscience, under review.

3. Barreiros, I. V, Panayi, M. C., & Walton, M. E. (2020). Organisation of afferents along the anterior-posterior and medial-lateral axes of the rat OFC. BioRxiv, 1–44. https://doi.org/https://doi.org/10.1101/2020.08.28.272591

4. Baxter, M. G., Parker, A., Lindner, C. C., Izquierdo, A. D., & Murray, E. A. (2000). Control of response selection by reinforcer value requires interaction of amygdala and orbital prefrontal cortex. Journal of Neuroscience, 20(11), 4311–4319. http://www.jneurosci.org/content/20/11/4311.full.pdf

5. Boulougouris, V., Dalley, J. W., & Robbins, T. W. (2007). Effects of orbitofrontal, infralimbic and prelimbic cortical lesions on serial spatial reversal learning in the rat. Behavioural Brain Research, 179(2), 219–228. https://doi.org/10.1016/j.bbr.2007.02.005

6. Boulougouris, V., & Robbins, T. W. (2009). Pre-surgical training ameliorates orbitofrontal-mediated impairments in spatial reversal learning. Behavioural Brain Research, 197(2), 469–475. https://doi.org/10.1016/j.bbr.2008.10.005

7. Bradfield, L. A., Dezfouli, A., Van Holstein, M., Chieng, B., & Balleine, B. W. (2015). Medial Orbitofrontal Cortex Mediates Outcome Retrieval in Partially Observable Task Situations. Neuron, 88(6), 1268–1280. https://doi.org/10.1016/j.neuron.2015.10.044

8. Bradfield, L. A., & Hart, G. (2020). Rodent medial and lateral orbitofrontal cortices represent unique components of cognitive maps of task space. Neuroscience & Biobehavioral Reviews, 108, 287–294. https://doi.org/10.1016/J.NEUBIOREV.2019.11.009

9. Bradfield, L. A., Hart, G., & Balleine, B. W. (2018). Inferring action-dependent outcome representations depends on anterior but not posterior medial orbitofrontal cortex. Neurobiology of Learning and Memory, 155(May), 463–473. https://doi.org/10.1016/j.nlm.2018.09.008

10. Burke, K. A., Franz, T. M., Miller, D. N., & Schoenbaum, G. (2007). Conditioned reinforcement can be mediated by either outcome-specific or general affective representations. Frontiers in Integrative Neuroscience, 1, 2. https://doi.org/10.3389/neuro.07.002.2007

11. Burke, K. A., Franz, T. M., Miller, D. N., & Schoenbaum, G. (2008). The role of the orbitofrontal cortex in the pursuit of happiness and more specific rewards. Nature, 454(7202), 340–U45. https://doi.org/Doi 10.1038/Nature06993

12. Burke, K. A., Takahashi, Y. K., Correll, J., Brown, P. L., & Schoenbaum, G. (2009). Orbitofrontal inactivation impairs reversal of Pavlovian learning by interfering with “disinhibition” of responding for previously unrewarded cues. European Journal of Neuroscience, 30(10), 1941–1946. https://doi.org/DOI 10.1111/j.1460-9568.2009.06992.x

13. Butter, C. M. (1969). Perseveration in extinction and in discrimination reversal tasks following selective frontal ablations in Macaca mulatta. Physiol. Behav, 4, 163–171.

14. Chudasama, Y., & Robbins, T. W. (2003). Dissociable contributions of the orbitofrontal and infralimbic cortex to pavlovian autoshaping and discrimination reversal learning: further evidence for the functional heterogeneity of the rodent frontal cortex. Journal of Neuroscience, 23(25), 8771–8780. https://doi.org/23/25/8771 [pii]

15. Collins, A. G. E., & Cockburn, J. (2020). Beyond dichotomies in reinforcement learning. Nature Reviews Neuroscience. https://doi.org/10.1038/s41583-020-0355-6

16. Coutureau, E., & Killcross, A. S. (2003). Inactivation of the infralimbic prefrontal cortex reinstates goal-directed responding in overtrained rats. Behavioural Brain Research, 146(1– 2), 167–174. http://www.sciencedirect.com/science/article/pii/S0166432803003498

17. Delamater, A. R. (2004). Experimental extinction in Pavlovian conditioning: Behavioural and neuroscience perspectives. Quarterly Journal of Experimental Psychology Section B-Comparative and Physiological Psychology, 57(2), 97–132. https://doi.org/Doi 10.1080/02724990344000097

18. Delamater, A. R. (2007). The role of the orbitofrontal cortex in sensory-specific encoding of associations in Pavlovian and instrumental conditioning. In G. Schoenbaum, J. A. Gottfried, E. A. Murray, & S. J. Ramus (Eds.), Linking Affect to Action: Critical Contributions of the Orbitofrontal Cortex (Vol. 1121, pp. 152–173). Blackwell Publishing. https://doi.org/10.1196/annals.1401.030

19. Delamater, A. R., & Holland, P. C. (2008). The influence of CS-US interval on several different indices of learning in appetitive conditioning. Journal of Experimental Psychology-Animal Behavior Processes, 34(2), 202–222. https://doi.org/Doi 10.1037/0097-7403.34.2.202

20. Delamater, A. R., & Oakeshott, S. (2007). Learning about multiple attributes of reward in Pavlovian conditioning. Annals of the New York Academy of Sciences. https://doi.org/10.1196/annals.1390.008

21. Dias, R., Robbins, T. W., & Roberts, A. C. (1996). Dissociation in prefrontal cortex of affective and attentional shifts. Nature, 380(6569), 69–72. https://doi.org/10.1038/380069a0

22. Dickinson, A. (1980). Contemporary animal learning theory. In J. Gray (Ed.), Problems in the behavioural sciences. Cambridge University Press.

23. Dickinson, A. (1985). Actions and Habits : The Development of Behavioural Autonomy. Philosophical Transactions of the Royal Society of London. Series B, Biological Sciences, 308(1135), 67–78. http://www.jstor.org/stable/2396284

24. Dickinson, A., & Balleine, B. W. (2002). The Role of Learning in the Operation of Motivational Systems. In Stevens’ Handbook of Experimental Psychology. John Wiley & Sons, Inc. https://doi.org/10.1002/0471214426.pas0312

25. Dolan, R. J., & Dayan, P. (2013). Goals and habits in the brain. Neuron, 80(2), 312–325. https://doi.org/10.1016/j.neuron.2013.09.007

26. Esber, G. R., & Haselgrove, M. (2011). Reconciling the influence of predictiveness and uncertainty on stimulus salience: a model of attention in associative learning. Proceedings of the Royal Society B-Biological Sciences, 278(1718), 2553–2561. https://doi.org/DOI 10.1098/rspb.2011.0836

27. Gallagher, M., McMahan, R. W., & Schoenbaum, G. (1999). Orbitofrontal cortex and representation of incentive value in associative learning. Journal of Neuroscience, 19(15), 6610–6614. http://www.jneurosci.org/cgi/reprint/19/15/6610.pdf

28. Gardner, M. P. H., Conroy, J. C., Sanchez, D. C., Zhou, J., & Schoenbaum, G. (2019). Real-Time Value Integration during Economic Choice Is Regulated by Orbitofrontal Cortex. Current Biology. https://doi.org/10.1016/j.cub.2019.10.058

29. Gardner, M. P. H., Conroy, J. C., Styer, C. V, Huynh, T., Whitaker, L. R., & Schoenbaum, G. (2018). Medial orbitofrontal inactivation does not affect economic choice. ELife, 7. https://doi.org/10.7554/eLife.38963

30. Gardner, M. P. H., Conroy, J. S., Shaham, M. H., Styer, C. V., & Schoenbaum, G. (2017). Lateral Orbitofrontal Inactivation Dissociates Devaluation-Sensitive Behavior and Economic Choice. Neuron, 0(0). https://doi.org/10.1016/j.neuron.2017.10.026

31. Hall, G. (2002). Associative structures in Pavlovian and instrumental conditioning. In C. R. Gallistel (Ed.), Steven’s handbook of experimental psychology (Vol. 3, pp. 1–45). John Wiley & Sons.

32. Holland, P. C. (1977). Conditioned stimulus as a determinant of the form of the Pavlovian conditioned response. J Exp Psychol Anim Behav Process, 3(1), 77–104. http://www.ncbi.nlm.nih.gov/pubmed/845545

33. Iversen, S. D., & Mishkin, M. (1970). Perseverative interference in monkeys following selective lesions of the inferior prefrontal convexity. Experimental Brain Research, 11(4), 376–386. http://www.ncbi.nlm.nih.gov/pubmed/4993199

34. Izquierdo, A. (2017). Functional Heterogeneity within Rat Orbitofrontal Cortex in Reward Learning and Decision Making. The Journal of Neuroscience : The Official Journal of the Society for Neuroscience, 37(44), 10529–10540. https://doi.org/10.1523/JNEUROSCI.1678-17.2017

35. Kamin, L. J. (1969). Predictability, surprise, attention and conditioning. In B. A. Campbell & R. M. Church (Eds.), Punishment and aversive behavior (pp. 279–96). Appleton-Century-Crofts.

36. Killcross, A. S., & Coutureau, E. (2003). Coordination of actions and habits in the medial prefrontal cortex of rats. Cerebral Cortex, 13(4), 400–408. http://cercor.oxfordjournals.org/cgi/reprint/13/4/400.pdf

37. Klein-Flugge, M. C., Barron, H. C., Brodersen, K. H., Dolan, R. J., & Behrens, T. E. (2013). Segregated encoding of reward-identity and stimulus-reward associations in human orbitofrontal cortex. Journal of Neuroscience, 33, 3202–3211. http://www.jneurosci.org/content/33/7/3202.full.pdf

38. Konorski, J. (1967). Integrative activity of the brain; an interdisciplinary approach. University of Chicago Press.

39. Kool, W., Cushman, F. A., & Gershman, S. J. (2018). Competition and cooperation between multiple reinforcement learning systems. In Goal-Directed Decision Making: Computations and Neural Circuits. Elsevier Inc. https://doi.org/10.1016/B978-0-12-812098-9.00007-3

40. Kringelbach, M. L. (2005). The human orbitofrontal cortex: Linking reward to hedonic experience. Nature Reviews Neuroscience, 6(9), 691–702. https://doi.org/Doi 10.1038/Nrn1747

41. Lasseter, H. C., Ramirez, D. R., Xie, X., & Fuchs, R. A. (2009). Involvement of the lateral orbitofrontal cortex in drug context-induced reinstatement of cocaine-seeking behavior in rats. European Journal of Neuroscience, 30(7), 1370–1381. https://doi.org/10.1111/j.1460-9568.2009.06906.x

42. Lenth, R., Singmann, H., Love, J., Buerkner, P., & Herve, M. (2020). emmeans: estimated marginal means. R package version 1.4. 4. In The American Statistician. https://doi.org/10.1080/00031305.1980.10483031

43. LePelley, M. E. (2004). The role of associative history in models of associative learning: A selective review and a hybrid model. Quarterly Journal of Experimental Psychology, 57B, 192–243.

44. Mackintosh, N. J. (1974). The psychology of animal learning. Academic Press.

45. Mackintosh, N. J. (1975). A theory of attention: Variations in the associability of stimuli with reinforcement. Psychol Rev, 82(4), 279–298. https://doi.org/10.1037/h0076778

46. McDannald, M. A., Lucantonio, F., Burke, K. A., Niv, Y., & Schoenbaum, G. (2011). Ventral striatum and orbitofrontal cortex are both required for model-based, but not model-free, reinforcement learning. Journal of Neuroscience, 31, 2700–2705. http://www.jneurosci.org/content/31/7/2700.full.pdf

47. McDannald, M. A., Saddoris, M. P., Gallagher, M., & Holland, P. C. (2005). Lesions of orbitofrontal cortex impair rats’ differential outcome expectancy learning but not conditioned stimulus-potentiated feeding. Journal of Neuroscience, 25(18), 4626–4632. https://doi.org/25/18/4626 [pii] 10.1523/JNEUROSCI.5301-04.2005

48. Murray, E. A., O’Doherty, J. P., & Schoenbaum, G. (2007). What we know and do not know about the functions of the orbitofrontal cortex after 20 years of cross-species studies. Journal of Neuroscience, 27(31), 8166–8169. https://doi.org/10.1523/JNEUROSCI.1556-07.2007

49. Murray, E. A., & Rudebeck, P. H. (2018). Specializations for reward-guided decision-making in the primate ventral prefrontal cortex. Nature Reviews Neuroscience, 19(7), 404–417. https://doi.org/10.1038/s41583-018-0013-4

50. Namboodiri, V. M. K., Otis, J. M., van Heeswijk, K., Voets, E. S., Alghorazi, R. A., Rodriguez-Romaguera, J., Mihalas, S., & Stuber, G. D. (2019). Single-cell activity tracking reveals that orbitofrontal neurons acquire and maintain a long-term memory to guide behavioral adaptation. Nature Neuroscience, 1. https://doi.org/10.1038/s41593-019-0408-1

51. Nasser, H. M., Calu, D. J., Schoenbaum, G., & Sharpe, M. J. (2017). The Dopamine Prediction Error: Contributions to Associative Models of Reward Learning. Frontiers in Psychology, 8, 244. https://doi.org/10.3389/fpsyg.2017.00244

52. Niv, Y. (2019). Learning task-state representations. Nature Neuroscience, 22(10), 1544–1553. https://doi.org/10.1038/s41593-019-0470-8

53. Ogawa, M., van der Meer, M. A. A., Esber, G. R., Cerri, D. H., Stalnaker, T. A., & Schoenbaum, G. (2013). Risk-responsive orbitofrontal neurons track acquired salience. Neuron, 77(2), 251–258. https://doi.org/10.1016/j.neuron.2012.11.006

54. Ostlund, S. B., & Balleine, B. W. (2007). Orbitofrontal cortex mediates outcome encoding in pavlovian but not instrumental conditioning. Journal of Neuroscience, 27(18), 4819–4825. https://doi.org/Doi 10.1523/Jneurosci.5443-06.2007

55. Padoa-Schioppa, C. (2009). Range-adapting representation of economic value in the orbitofrontal cortex. Journal of Neuroscience, 29, 14004–14014. http://www.jneurosci.org/content/29/44/14004.full.pdf

56. Panayi, M. C., & Killcross, S. (2018). Functional heterogeneity within the rodent lateral orbitofrontal cortex dissociates outcome devaluation and reversal learning deficits. ELife, 7. https://doi.org/10.7554/eLife.37357.001

57. Pearce, J. M., & Hall, G. (1980). A model for Pavlovian learning: variations in the effectiveness of conditioned but not of unconditioned stimuli. Psychol Rev, 87(6), 532–552. http://www.ncbi.nlm.nih.gov/pubmed/7443916

58. Pears, A., Parkinson, J. A., Hopewell, L., Everitt, B. J., & Roberts, A. C. (2003). Lesions of the orbitofrontal but not medial prefrontal cortex disrupt conditioned reinforcement in primates. Journal of Neuroscience, 23(35), 11189–11201. https://doi.org/23/35/11189 [pii]

59. Pickens, C. L., Saddoris, M. P., Gallagher, M., & Holland, P. C. (2005). Orbitofrontal lesions impair use of cue-outcome associations in a devaluation task. Behav Neurosci, 119(1), 317– 322. https://doi.org/2005-01705-030 [pii] 10.1037/0735-7044.119.1.317

60. Pickens, C. L., Saddoris, M. P., Setlow, B., Gallagher, M., Holland, P. C., & Schoenbaum, G. (2003). Different Roles for Orbitofrontal Cortex and Basolateral Amygdala in a Reinforcer Devaluation Task. The Journal of Neuroscience, 23(35), 11078–11084. https://doi.org/10.1523/JNEUROSCI.23-35-11078.2003

61. R Core Team (2020). (2020). R: A language and environment for statistical computing. In R: A language and environment for statistical computing. *R Foundation for Statistical Computing*, *Vienna, Austria*.

62. Ramirez, D. R., & Savage, L. M. (2007). Differential involvement of the basolateral amygdala, orbitofrontal cortex, and nucleus accumbens core in the acquisition and use of reward expectancies. Behav Neurosci, 121(5), 896–906. https://doi.org/10.1037/0735-7044.121.5.896

63. Rescorla, R. A. (1988). Pavlovian Conditioning - Its Not What You Think It Is. American Psychologist, 43(3), 151–160. https://doi.org/Doi 10.1037/0003-066x.43.3.151

64. Rescorla, R. A. (2002a). Comparison of the rates of associative change during acquisition and extinction. Journal of Experimental Psychology: Animal Behavior Processes, 28(4), 406– 415. https://doi.org/10.1037/0097-7403.28.4.406

65. Rescorla, R. A. (2002b). Savings tests: Separating differences in rate of learning from differences in initial levels. Journal of Experimental Psychology: Animal Behavior Processes, 28(4), 369–377. https://doi.org/10.1037/0097-7403.28.4.369

66. Rescorla, R. A., & Wagner, A. R. (1972). A theory of Pavlovian conditiong: Variations in the effectiveness of reinforcement and nonreinforcement. In A. H. Black & W. F. Prokesy (Eds.), Classical Conditioning II: Current Research and Theory (pp. 64–99). Appleton Century Crofts.

67. Rudebeck, P. H., & Murray, E. A. (2014). The Orbitofrontal Oracle: Cortical Mechanisms for the Prediction and Evaluation of Specific Behavioral Outcomes. Neuron, 84(6), 1143–1156. https://doi.org/10.1016/j.neuron.2014.10.049

68. Sadacca, B. F., Wied, H. M., Lopatina, N., Saini, G. K., Nemirovsky, D., & Schoenbaum, G. (2018). Orbitofrontal neurons signal sensory associations underlying model-based inference in a sensory preconditioning task. ELife, 7, e30373. https://doi.org/10.7554/eLife.30373

69. Schoenbaum, G., Nugent, S. L., Saddoris, M. P., & Setlow, B. (2002). Orbitofrontal lesions in rats impair reversal but not acquisition of go, no-go odor discriminations. Neuroreport, 13(6), 885–890. https://doi.org/10.1097/00001756-200205070-00030

70. Schoenbaum, G., Roesch, M. R., Stalnaker, T. A., & Takahashi, Y. K. (2009). A new perspective on the role of the orbitofrontal cortex in adaptive behaviour. Nature Reviews Neuroscience, 10(12), 885–892. https://doi.org/Doi 10.1038/Nrn2753

71. Schoenbaum, G., Setlow, B., Nugent, S. L., Saddoris, M. P., & Gallagher, M. (2003). Lesions of orbitofrontal cortex and basolateral amygdala complex disrupt acquisition of odor-guided discriminations and reversals. Learning & Memory, 10(2), 129–140. https://doi.org/Doi 10.1101/Lm.55203

72. Schoenbaum, G., Takahashi, Y. K., Liu, T., & McDannald, M. A. (2011). Does the orbitofrontal cortex signal value? Ann N Y Acad Sci, 1239(1), 87–99. https://doi.org/10.1111/j.1749-6632.2011.06210.x

73. Schultz, W. (1998). Predictive reward signal of dopamine neurons. Journal of Neurophysiology, 80(1), 1–27.

74. Sharpe, M. J., Chang, C. Y., Liu, M. A., Batchelor, H. M., Mueller, L. E., Jones, J. L., Niv, Y., & Schoenbaum, G. (2017a). Dopamine transients are sufficient and necessary for acquisition of model-based associations. Nature Neuroscience. https://doi.org/10.1038/nn.4538

75. Sharpe, M. J., Chang, C. Y., Liu, M. A., Batchelor, H. M., Mueller, L. E., Jones, J. L., Niv, Y., & Schoenbaum, G. (2017b). Dopamine transients are sufficient and necessary for acquisition of model-based associations. Nature Neuroscience, 20(5), 735–742. https://doi.org/10.1038/nn.4538

76. Sharpe, M. J., Wikenheiser, A. M., Niv, Y., & Schoenbaum, G. (2015). The State of the Orbitofrontal Cortex. Neuron, 88(6), 1075–1077. https://doi.org/10.1016/j.neuron.2015.12.004

77. Singmann, H., Bolker, B., Westfall, J., Aust, F., Højsgaard, S., Fox, J., Lawrence, M. A., Mertens, U., & Love, J. (2020). afex: analysis of factorial experiments. R Package Version 0.27-2. https://cran.r-project.org/package=afex

78. Stalnaker, T. A., Cooch, N. K., McDannald, M. A., Liu, T. L., Wied, H., Schoenbaum, G., & Tzu-Lan, L. (2014). Orbitofrontal neurons infer the value and identity of predicted outcomes. Nat Commun, 5, 3926. https://doi.org/10.1038/ncomms4926

79. Stalnaker, T. A., Cooch, N. K., & Schoenbaum, G. (2015). What the orbitofrontal cortex does not do. In Nature Neuroscience. https://doi.org/10.1038/nn.3982

80. Stalnaker, T. A., Franz, T. M., Singh, T., & Schoenbaum, G. (2007). Basolateral amygdala lesions abolish orbitofrontal-dependent reversal impairments. Neuron, 54(1), 51–58. https://doi.org/DOI 10.1016/j.neuron.2007.02.014

81. Stalnaker, T. A., Liu, T.-L., Takahashi, Y. K., & Schoenbaum, G. (2018). Orbitofrontal neurons signal reward predictions, not reward prediction errors. Neurobiology of Learning and Memory, 153, 137–143. https://doi.org/10.1016/J.NLM.2018.01.013

82. Steinberg, E. E., Keiflin, R., Boivin, J. R., Witten, I. B., Deisseroth, K., & Janak, P. H. (2013). A causal link between prediction errors, dopamine neurons and learning. Nature Neuroscience, 16(7), 966–973. https://doi.org/10.1038/nn.3413

83. Stolyarova, A., & Izquierdo, A. (2017). Complementary contributions of basolateral amygdala and orbitofrontal cortex to value learning under uncertainty. ELife, 6, e27483. https://doi.org/10.7554/eLife.27483

84. Sutton, R. S., & Barto, A. G. (1998). Reinforccement Learning: An introduction. The MIT Press.

85. Takahashi, Y. K., Chang, C. Y., Lucantonio, F., Haney, R. Z., Berg, B. A., Yau, H.-J., Bonci, A., & Schoenbaum, G. (2013). Neural estimates of imagined outcomes in the orbitofrontal cortex drive behavior and learning. Neuron, 80, 507–518. http://ac.els-cdn.com/S0896627313007198/1-s2.0-S0896627313007198-main.pdf?_tid=97ea45dc-cbcc-11e4-9c0e-00000aacb362&acdnat=1426504211_80b43da207445d70382dd3274f8c4f11

86. Takahashi, Y. K., Roesch, M. R., Stalnaker, T. A., Haney, R. Z., Caiu, D. J., Taylor, A. R., Burke, K. A., Schoenbaum, G., & Calu, D. J. (2009). The Orbitofrontal Cortex and Ventral Tegmental Area Are Necessary for Learning from Unexpected Outcomes. Neuron, 62(2), 269–280. https://doi.org/DOI 10.1016/j.neuron.2009.03.005

87. Takahashi, Y. K., Roesch, M. R., Wilson, R. C., Toreson, K., O’Donnell, P., Niv, Y., & Schoenbaum, G. (2011). Expectancy-related changes in firing of dopamine neurons depend on orbitofrontal cortex. Nature Neuroscience, 14(12), 1590–1597. https://doi.org/10.1038/nn.2957

88. Wagner, A. R. (1981). SOP: A model of automatic memory processing in animal behavior. In N. E. Spear (Ed.), Information processing in animals: Memory mechanisms (pp. 5–47). Erlbaum.

89. Walton, M. E., Behrens, T. E., Buckley, M. J., Rudebeck, P. H., & Rushworth, M. F. (2010). Separable learning systems in the macaque brain and the role of orbitofrontal cortex in contingent learning. Neuron, 65(6), 927–939. https://doi.org/10.1016/j.neuron.2010.02.027

90. Walton, M. E., Behrens, T. E., Noonan, M. P., & Rushworth, M. F. (2011). Giving credit where credit is due: orbitofrontal cortex and valuation in an uncertain world. Ann N Y Acad Sci, 1239, 14–24. http://onlinelibrary.wiley.com/doi/10.1111/j.1749-6632.2011.06257.x/abstract

91. Wang, P. Y., Boboila, C., Chin, M., Higashi-Howard, A., Shamash, P., Wu, Z., Stein, N. P., Abbott, L. F., & Axel, R. (2020). Transient and Persistent Representations of Odor Value in Prefrontal Cortex. Neuron, 1–16. https://doi.org/10.1016/j.neuron.2020.07.033

92. West, E. A., DesJardin, J. T., Gale, K., & Malkova, L. (2011). Transient Inactivation of Orbitofrontal Cortex Blocks Reinforcer Devaluation in Macaques. The Journal of Neuroscience : The Official Journal of the Society for Neuroscience, 31(42), 15128–15135. https://doi.org/10.1523/JNEUROSCI.3295-11.2011

93. Wilson, R. C., Takahashi, Y. K., Schoenbaum, G., & Niv, Y. (2014). Orbitofrontal cortex as a cognitive map of task space. Neuron, 81(2), 267–279. https://doi.org/10.1016/j.neuron.2013.11.005

94. Zhou, J., Gardner, M. P. H., Stalnaker, T. A., Ramus, S. J., Wikenheiser, A. M., Niv, Y., & Schoenbaum, G. (2019). Rat Orbitofrontal Ensemble Activity Contains Multiplexed but Dissociable Representations of Value and Task Structure in an Odor Sequence Task. Current Biology. https://doi.org/10.1016/j.cub.2019.01.048

